# Adiposome Targeting and Enzymatic Activity of Lipid Droplet-Specific Proteins

**DOI:** 10.1101/2020.04.27.062869

**Authors:** Xuejing Ma, Zelun Zhi, Shuyan Zhang, Chang Zhou, Adam Mechler, Pingsheng Liu

## Abstract

New strategies to decode the specific protein targeting mechanism on lipid droplet (LD) are urgently needed. Using adiposome, the LD binding of perilipin 2 (PLIN2), perilipin 3 (PLIN3), and adipose triglyceride lipase (ATGL) were studied. Scatchard analysis found that the binding of PLIN2 to the adiposome surface was saturable, pointing to a specific membrane binding partner. Phosphatidylinositol (PI) was found to inhibit PLIN2 binding while it did not impede PLIN3. Structural analysis combined with mutagenesis revealed that the 73^rd^ glutamic acid of PLIN2 is significant for the effect of PI on the protein binding. The presence of PI significantly stimulated the activity of ATGL *in vitro*. The phosphorylation site mutants of ATGL were found reducing the lipase activity in the adiposome system. Our study demonstrates the utility of adiposome as a powerful, manipulatable model system for the characterization of LD binding and enzymatic activity of LD proteins *in vitro*.

## INTRODUCTION

The lipid droplet (LD) is a unique organelle conserved in most organisms from bacteria to humans (Martin and Parton, 2006; Zhang and Liu, 2017). The LD consists of a neutral lipid core of mostly triacylglycerol (TAG), sterol esters, retinyl ester, and/or polyhydroxyalkanoate, surrounded by a monolayer phospholipid membrane and associated proteins (Farese and Walther, 2009). While an understanding of the functions of the LD as an organelle is still developing, it is certain that it plays a significant role in energy homeostasis, in particular lipid metabolism, storage, and transportation (Bartz et al., 2007a; Liu et al., 2004; Walther et al., 2017; Yao et al., 2019). Proteomic and lipidomic studies using isolated LDs have provided detailed information on the protein and lipid composition of the organelle from almost all model organisms (Liu et al., 2004; Zhang and Liu, 2019). Many of the LD proteins identified through proteomics have also been studied in morphological and functional studies (Ding et al., 2012; Liu et al., 2004; Na et al., 2015). Most LD-associated proteins are found at multiple locations in cells. However, a smaller number are resident proteins that specifically and directly localize on the organelle (Na et al., 2015; Ohsaki et al., 2014; Wolins et al., 2006; Zhang and Liu, 2019). With few exceptions these proteins are only found on the LD surface, and therefore it is believed that they possess the binding mechanism(s) specific for monolayer phospholipid membranes, the nature of which is under debate.

Several studies have attempted to identify targeting mechanisms with limited success (Walther and Farese, 2012). Based on these findings targeting mechanisms can be roughly divided into two groups, direct and indirect (Zhang and Liu, 2019). Indirect localization is mainly maintained by acylation and protein-protein interaction (Leung et al., 2007; Shen et al., 2009). For direct localization, specific domains have been identified that are integral monotopic structures including hydrophobic domains or/and amphipathic α-helixes (Bersuker and Olzmann, 2017; Kory et al., 2016). The targeting formats include hydrophobic hairpin, terminal hydrophobic domain, and non-terminal hydrophobic domains (Boeszoermenyi et al., 2015; Huang and Huang, 2017; Na et al., 2015). In addition, one or more amphipathic α-helixes could be also involved (Ding et al., 2012; Krahmer et al., 2011), a type of motif that is characteristic of LD resident proteins (Bulankina et al., 2009; Chong et al., 2011; Rowe et al., 2016; Subramanian et al., 2004). Targeting domains have been identified using protein truncation mutants (Nakamura and Fujimoto, 2003). A computer simulation has also been used to direct biochemical assays and to suggest a possible binding mechanism (Prévost et al., 2018). Regardless, no definitive solution to the mechanism(s) which permits selective LD binding has been determined. *In vivo* assays may be insufficient to dissect the binding mechanism due to the complexity of the cellular environment. Therefore, it is crucial to develop an *in vitro* method to simplify binding conditions and combine *in vitro* and *in vivo* assays to provide detailed insights into the targeting mechanism.

Adiposomes are artificial nanostructures containing a neutral lipid core coated with a monolayer phospholipid membrane, which can be used to mimic the structure and function of LDs to allow *in vitro* LD assays (Wang et al., 2016; Zhang et al., 2017). Compared to previous methods used to prepare LD-like emulsions, our technique separates adiposomes from impurities and keeps the diameter of adiposomes homogeneously, allowing the structure to more effectively model LDs (Chen et al., 2015; Fei et al., 2011; Krahmer et al., 2011; Tzen and Huang, 1992; Wang et al., 2016). Importantly, the neutral and polar lipid constituents of adiposomes can be controlled as required for a particular experiment. For example, the role of protein-phospholipid binding in membrane targeting can be explored through manipulation of the adiposome phospholipid composition (Lemmon, 2008; Yan et al., 2018). The role of phosphatidic acid in LD protein targeting has been reported (Barneda et al., 2015; Yan et al., 2018), but the role of phosphatidyl inositol (PI), which is abundant on the LD, has attracted little attention (Bartz et al., 2007a; Tauchi-Sato et al., 2002). However, PI is known to be involved in protein binding in other contexts (Phan et al., 2016), so further investigation is well warranted.

The LD is a reservoir of neutral lipids, especially TAG, and also is a site of lipid synthesis and lipolysis (Walther and Farese, 2012). Therefore, a change in lipase activity on LDs will affect cellular lipid metabolism and homeostasis. Adipose triglyceride lipase (ATGL) is a TAG hydrolytic enzyme with multiple identified phosphorylation sites (Ahmadian et al., 2011; Bartz et al., 2007b; Pagnon et al., 2012; Xie et al., 2014). Active ATGL targeted to LDs can reduce their volume, even to the point where they become undetectable, making the study of ATGL targeting challenging. For this reason, adiposome provide an ideal model system to study the phosphorylation dependent targeting and regulation of ATGL.

To study LD protein targeting mechanisms, we constructed an adiposome system based on our previous work. The binding affinity of the LD resident PLIN2 to the monolayer phospholipid membrane was studied with an emphasis on the role of the anionic phospholipid phosphatidylinositol (PI). The targeting mechanism of PLIN2 was also assessed along with the binding and activity of ATGL. Our results demonstrate that the adiposomes closely mimic the actual LDs for *in vitro* studies of protein targeting and activity.

## RUSULTS

### Construction of protein binding adiposome *in vitro* platform

The preparation of adiposome followed the method reported by our group previously (Wang et al., 2016). To verify the availability of adiposome on mimicking the protein binding to LD, the binding of GFP-tagged PLIN2 to natural LDs and adiposomes was compared morphologically and quantitatively. A PLIN2-GFP knock-in C2C12 cell line was constructed to study the distribution of PLIN2 *in vivo*, as recently reported (Xu et al., 2019). To evaluate binding to adiposomes in an *in vitro* system we expressed and purified recombinant SMT3-PLIN2-GFP and PLIN3-APPLE. The purified proteins were analyzed by Colloidal Blue staining and Western blot (anti-PLIN2 antibody for SMT3-PLIN2-GFP and anti-PLIN3 antibody for PLIN3-APPLE, Abcam) (Figure S1). The rough purities of SMT3-PLIN-GFP and PLIN3-GFP, as determined using ImageJ, were 59% and 86%, respectively. The proteins were mixed with adiposomes constructed of DOPC and TAG. Both GFP-tagged PLIN2 fusion proteins appeared as ring-like structures around LDs (Figure 1A) and adiposomes (Figure 1B), confirming that both the endogenous and recombinant PLIN2 were able to target to LDs and adiposomes, respectively.

**Figure 1.**
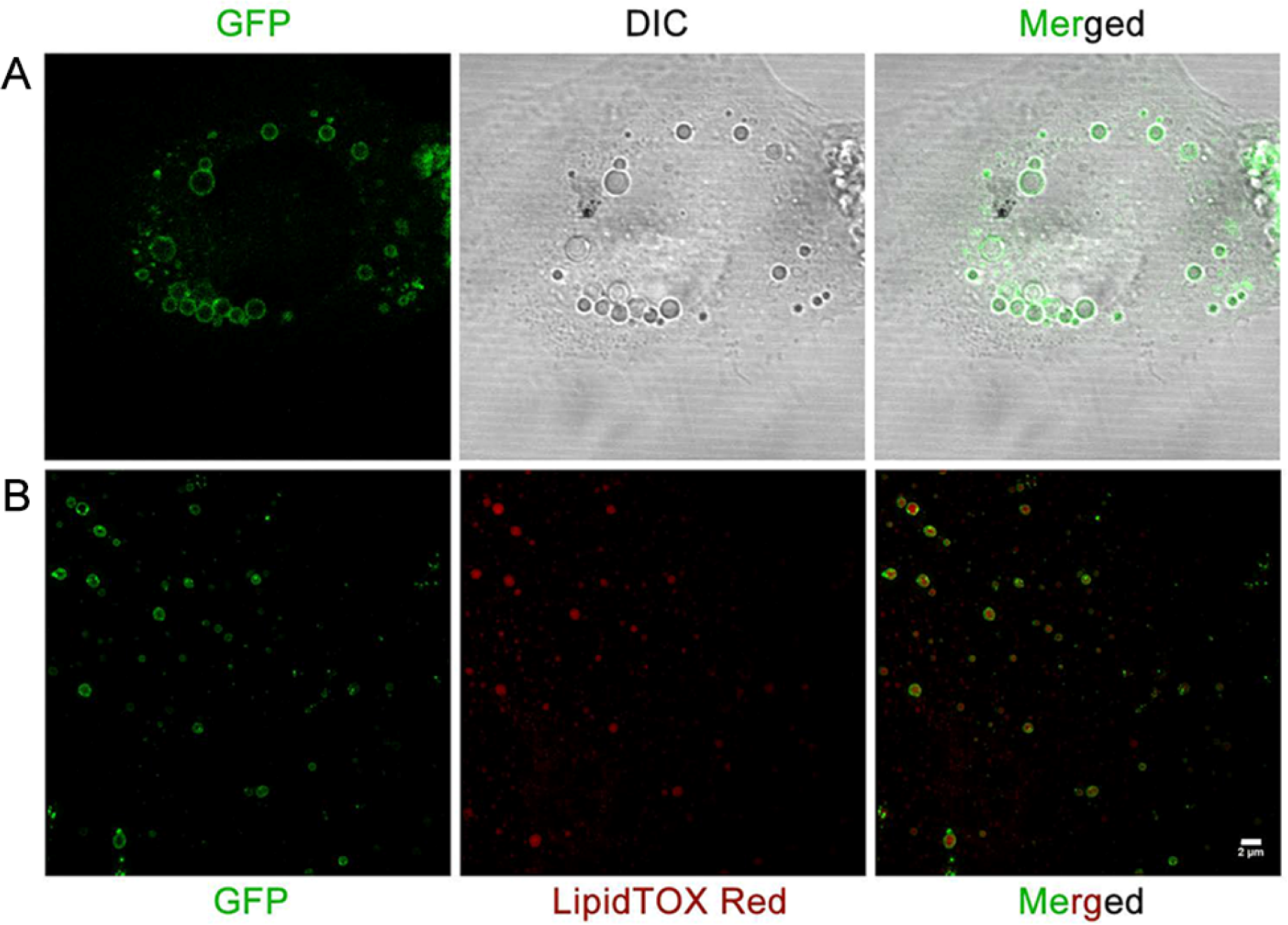
PLIN2 targets to LDs and adiposomes. (A) PLIN2–GFP knock-in C2C12 cells were seeded in confocal dishes overnight. After 100 μM oleate treatment for 12 h, cells were imaged by Olympus FV1000 confocal microscope. Lipid droplets became much larger due to OA treatment and PLIN2-GFP could target on lipid droplets forming green ringed structures. (B) The adiposomes prepared from neat DOPC and TAG were incubated with SMT3-PLIN2-GFP. Fluorescence images were captured using a DeltaVision OMX (SIM) microscope following LipidTOX Red staining. The fusion protein formed an even distribution over the adiposome surface. Scale bar, 2 μm.

We next determined the density of expressed PLIN2 on LDs and recombinant PLIN2 on adiposomes. First the concentration of a preparation of purified, recombinant PLIN2 was determined by comparison to BSA using densitometric analysis of Coomassie stained gels (Figure S2A). The density of PLIN2 on the adiposome was then calculated using the following the equation:

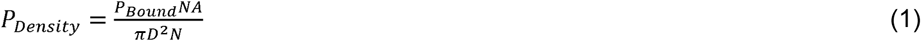

where D is the diameter of the LDs/adiposomes, P_Bound_ is the maximum amount of the protein on the surface of LDs/adiposomes, NA is Avogadro’s constant, and N is the absolute number of LDs/adiposomes. The concentration of PLIN2-GFP on LDs isolated from PLIN2-GFP knock-in (KI) cells was determined using a similar approach. The quantity of PLIN2-GFP on LDs was estimated by comparing the Western blot signal of the LDs to a standard curve of recombinant PLIN2 (Figure S2B) (Poppelreuther et al., 2018). The average LD diameter was determined to be approximately 540.2 nm using dynamic light scattering and the concentration of LDs was determined to be approximately 2.26 × 10^8^ per ml using FFF-MALS (Figure S2C and D). Multiplying the average surface area of a LD by the LD concentration gave a total surface area of 8.28 × 10^6^ μm^2^ per ml. The estimated concentration of PLIN2 on LDs was 7.73 × 10^14^ per ml, derived from the quantification of the Western blot. Hence, the density of endogenous GFP-tagged PLIN2 on LDs was 9.33 × 10^7^ per μm^2^. The average diameter of DOPC adiposomes was determined to be 161.1 nm using dynamic light scattering (Figure 2B) and the number of adiposomes was estimated as 8.53 × 10^9^ per ml by FFF-MALS (Figure S2E). The density of SMT3-PLIN2-GFP on the adiposome was calculated as 1.12 × 10^6^ per μm^2^. In both cases even distribution was observed in the fluorescence images that lead to a ring-like appearance. Figure S2F shows the morphological difference of adiposomes and lipid emulsions using TEM with ultrathin section. Clearly, lipid emulsions contained non-spherical multiplayer structures and LD-like droplets, while adiposomes were pure spherical LD-like droplets. The characterization of adiposome structure, phospholipid monolayer covering neutral lipid core, has been proved by our early report (Wang et al., 2016). Previous studies on *in vitro* LD-specific protein targeting usually used lipid emulsions, and bilayer structure like large unilamellar vesicles as the LD-mimicking model (Krahmer et al., 2011; Prévost et al., 2018; Sletten et al., 2014). Those models, either failed to mimic the monolayer and neutral lipid structure of LD, or did not show a homogeneous size distribution of droplets, making them suitable for fundamental protein binding research, but insufficient for precisely quantitative determination of LD protein binding. The adiposome model therefore shows advantages than those reported models.

**Figure 2.**
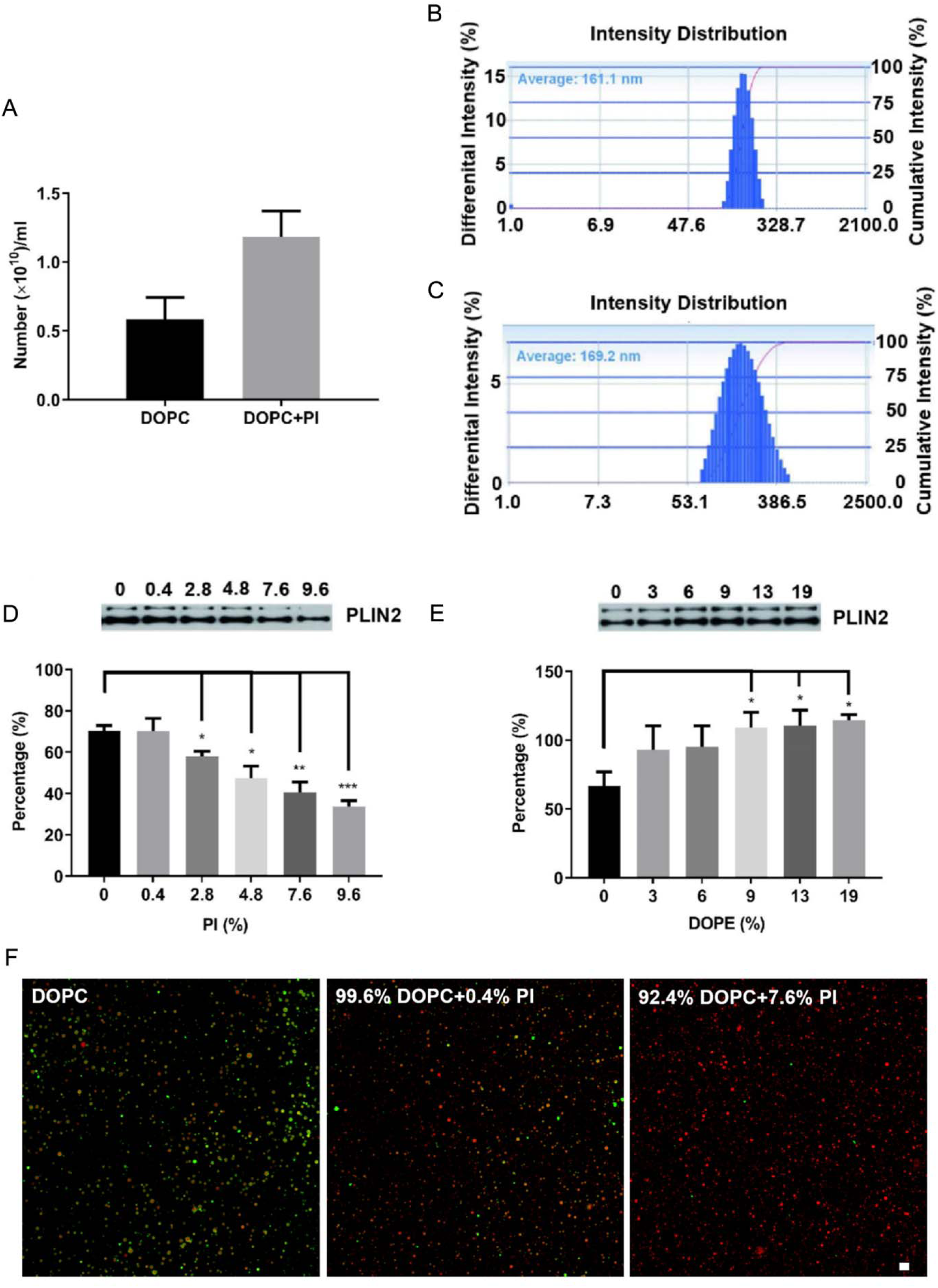
Impact of phospholipids composition on PLIN2 association to adiposomes. (A) The concentration of adiposomes with PI (92.4% DOPC + 7.6% PI) or without PI in a preparation with an OD600 = 20 (using Eppendorf Biophotometer) shows no significant difference (determined by FFF-MALS). Data represent mean ± s.e.m., n = 3. (B) The average diameter of adiposomes prepared with DOPC determined using DLS was 161.1 nm. Polydispersity Index = 0.121. (C) The average diameter of adiposomes prepared by DOPC (92.4%, mol/mol) and PI (7.6%, mol/mol) determined using DLS was 169.2 nm. Polydispersity Index = 0.177. (D) The targeting of PLIN2 on adiposomes was reduced when the ratio of PI increased, (E) while it was increased when the ratio of DOPE increased. All the Western blots were performed using anti-PLIN2 antibody. The grey scale was analyzed by ImageJ. Data represent mean ± s.e.m., n = 3. \**P* < 0.05, \*\**P* < 0.01, *\*\*\*P* < 0.001, two-tailed t-test. (F) The fluorescence images of PLIN2 (green) targeting on different adiposomes (red). Fluorescence was imaged using a Confocal FV1000 microscope following three washes and LipidTOX Red staining. Scale bar, 2 μm.

The range of PLIN2-GFP protein concentration giving a linear fluorescence response was determined (Figure S3A and B). The addition of Triton X-100 was required in the pure protein solution to prevent protein aggregation, which distorted the relationship. The relationship between adiposome quantity and OD600 is shown in Figure S3C. The experiments were performed with consideration of the non-linear response in OD600 at high concentrations of adiposomes (Figure S3D). In experiments performed in the presence of adiposomes, no Triton X-100 was required to maintain a linear fluorescence response with increasing PLIN2-GFP (Figure S3E). The data suggest that adiposomes facilitated the dispersion of protein aggregates. According to the results in Figure 3B, the saturated concentration was 0.59 μM (average value of three replicates) and the concentration of maximum bound SMT3-PLIN2-GFP was 1.29 μM. Hence, the density of the protein on adiposomes was 1.12 × 10^6^ per μm^2^. Compared to the density of PLIN2 on LDs, adiposomes are able to be used as LD-mimics for protein binding studies.

**Figure 3.**
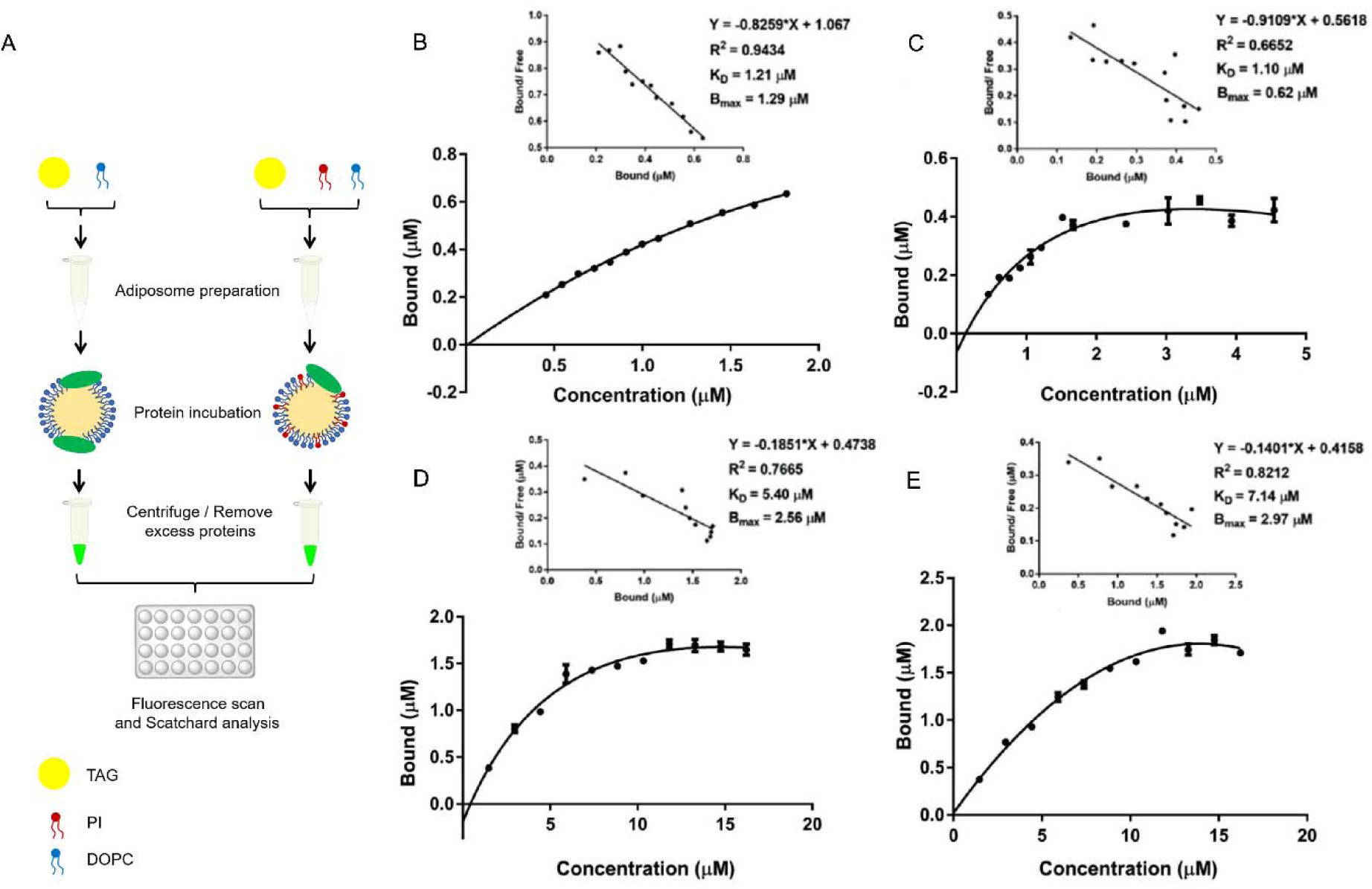
Scatchard analyses of PLIN2 and PLIN3 targeting to adiposomes. (A) Diagram of the experimental design. The saturation curves with Scatchard plot of SMT3-PLIN2-GFP on adiposomes (B) in the absence of PI or (C) in the presence of PI (92.4% DOPC + 7.6% PI) were measured using EnSpire Multimode Plate Reader and analyzed by GraphPad 7.0. The x-axis represents the concentration of PLIN2 in both Figures. The saturation of PLIN3-APPLE on adiposomes (D) without PI or (E) with PI (92.4% DOPC + 7.6% PI) was measured and analyzed using the same method for SMT3-PLIN2-GFP. The x-axis represents the concentration of PLIN3 in both Figures. The data were collected from three technical replicates for Scatchard analysis. The concentration of purified proteins was determined by BCA and the fluorescence intensity (FI) represents their concentrations on adiposomes. A series of concentrations of SMT3-PLIN2-GFP and PLIN3-APPLE were incubated with adiposomes at 4°C for 12 h and the FI was detected using EnSpire Multimode Plate Reader. PLIN2 recruitment to adiposomes was saturable. There was a significant difference of the saturation concentration of PLIN2 on adiposomes with or without PI, whereas there were no differences in saturation of PLIN3 on adiposomes with or without PI. The equilibrium dissociation constant K_d_ and the maximum saturation concentration of the binding sites on the adiposome, B_max_, were determined using Scatchard plots, (B) K_d(PLIN2)_ = 1.21 μM, B_max_ = 1.29 μM in the absence of PI, (C) K_d(PLIN2)_ = 1.10□μM, B_max_ = 0.62 μM in the presence of PI, (D) K_d(PLIN3)_ = 5.40 μM, B_max_ = 2.56 μM in the absence of PI and (E) K_d(PLIN3)_ = 7.14□μM, B_max_ = 2.97 μM in the presence of PI.

### Pl inhibits the targeting of PLIN2 to adiposomes

PI is the dominant negatively charged phospholipid on the LD monolayer membrane (Bartz et al., 2007a). We investigated whether PI plays a role in the targeting of PLIN2, which is one of the major resident proteins on LDs (Miura et al., 2002; Nakamura and Fujimoto, 2003). Adiposomes consisting of varied DOPC/PI ratios were constructed to investigate the role of PI in protein targeting. Adiposomes composed of DOPC and DOPE were used as a control for the targeting study. The physiological ratios of the primary membrane lipids are known: the molar ratios of PC, PE and PI to total phospholipids on LD are roughly 46%, 17% and 8%, respectively (Bartz et al., 2007a). The concentration (Figure 2A) and size distribution (Figure 2B and C) of adiposomes prepared with and without PI were very similar, guaranteeing a consistent adiposome quantity and identical light scattering behavior for each assay. Therefore, adiposome concentrations of the different preparations could be normalized using OD600. The OD600 values of the adiposome samples were measured before they were incubated with proteins to ensure equivalence. Western blot of PLIN2-incubated adiposomes showed that the targeting of PLIN2 to adiposomes increased in proportion to DOPE content, whereas it was gradually reduced with increasing PI content (Figure 2D and E). Fluorescence imaging confirmed this trend: Figure 2F shows that the green ringed structures formed by the recombinant PLIN2 gradually disappeared with increasing PI ratio, while the density of adiposomes (red dots) remained roughly constant. Thus, targeting of PLIN2 was inhibited by PI.

The presence of PI in the adiposome monolayer membrane inhibited the targeting of PLIN2. It is unknown if this is generalizable to other LD-associated proteins. Therefore, adiposome binding of PLIN3, another major LD-associated protein in Perilipin family, was compared with PLIN2. Recombinant PLIN3 with an APPLE tag was incubated with adiposomes with varying PI content using the same method as for PLIN2. Figure 3 summarizes the procedure and results of the binding experiments. The binding kinetic charts and Scatchard plots show that both PLIN2 and PLIN3 were able to bind to adiposomes in a saturable pattern. Based on the Scatchard analysis, the B_max_ value for SMT3-PLIN2-GFP binding to adiposome in the absence of PI (1.29 μM) was higher than that binding on the adiposome in the presence of PI (0.62 μM) (Figure 3B and C). This is consistent with the fluorescence microscopy results showing that PI reduced the binding of SMT3-PLIN2-GFP on adiposomes. In contrast, the B_max_ value of PLIN3-APPLE on DOPC adiposomes was 2.56 μM while on PI containing adiposome was 2.97 μM, overall higher values than those of SMT3-PLIN2-GFP (Figure 3D and E). The K_d_ values for SMT3-PLIN2-GFP either in the presence or absence of PI was lower than the values of PLIN3-APPLE, suggesting a higher affinity of SMT3-PLIN2-GFP for adiposomes than PLIN3-APPLE (K_d(PLIN2)_ = 1.21 μM, K_d(PLIN3)_ = 5.40 μM in the absence of PI; K_d(PLIN2)_ = 1.10□μM, K_d(PLIN3)_ = 7.14□μM in the presence of PI). This is consistent with the known difference between the two proteins: PLIN2 is an LD resident protein while PLIN3 is described as LD dynamic protein (Miura et al., 2002).

### The 73^rd^ glutamic acid in PLIN2 significantly affects the targeting of PLIN2

To investigate the mechanism of their differences in targeting, the structures of PLIN2 and PLIN3 were compared. The full native structures of PLIN2 and PLIN3 *(Homo sapiens)* have not yet been solved. Therefore, the known crystal structure of PLIN3 _(C-terminal)_ (*Mus musculus)* aa 191-437 (PDB ID: 1SZI) was used as a template to predict the structures of PLIN2 and PLIN3 (*Homo sapiens)* using I-TASSER. The structures with the highest confidence scores for the proteins were selected for structural comparison using PyMOL (Hickenbottom et al., 2004). Figure 4A shows the optimized structures of PLIN2 (blue) and PLIN3 (red). The secondary structure of PLIN2 and PLIN3 was predicted by Phyre2.0 and the sequence of PLIN2 was visualized by IBS (Figure S4) (Liu et al., 2015). The structurally different regions are marked in pink and the anionic amino acids in the amphiphilic helixes of PLIN2 are shown in yellow.

**Figure 4.**
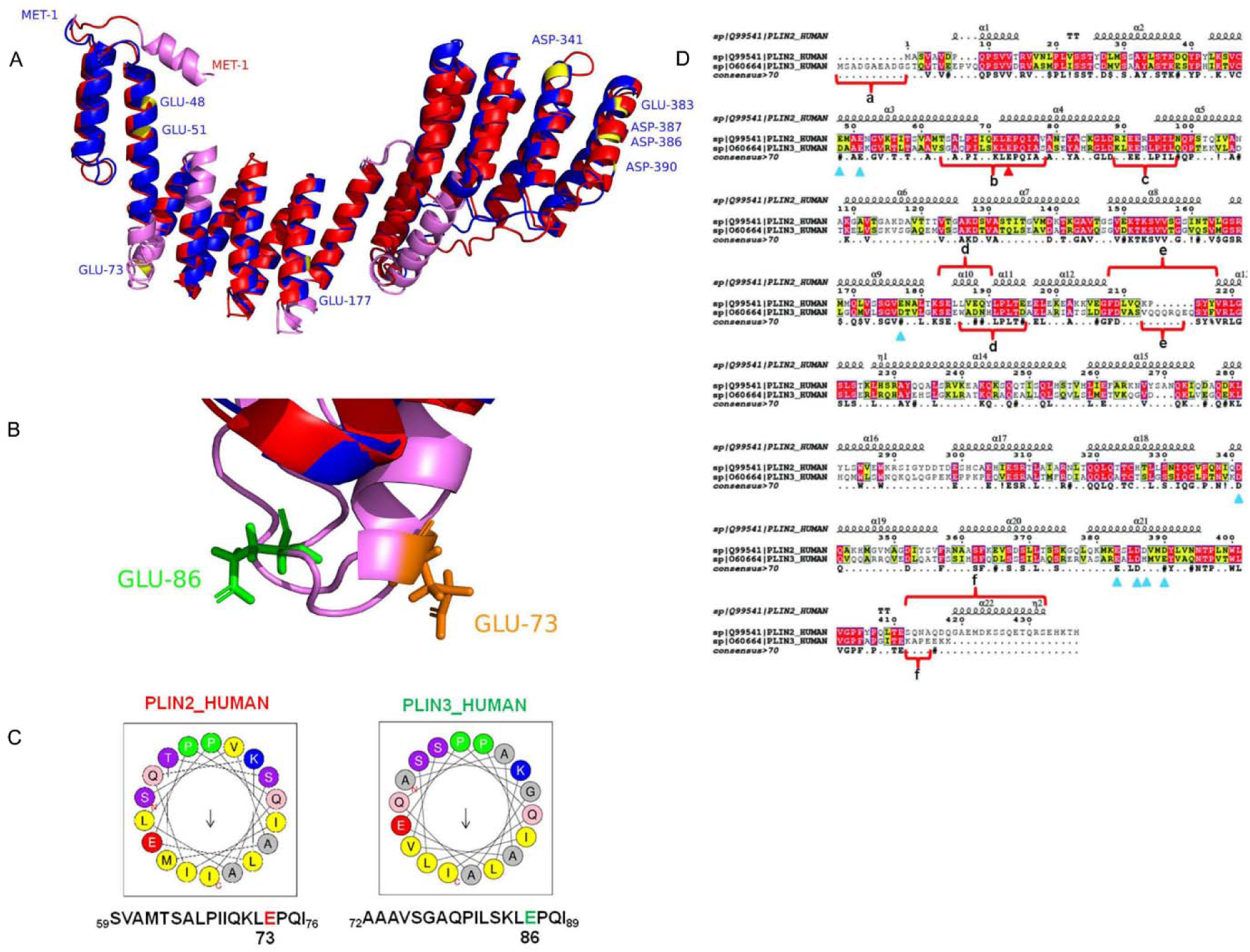
Structural analysis reveals the differences between PLIN2 and PLIN3. (A) The overall structural comparison of PLIN2 (blue) and PLIN3 (red) shows structural differences (pink) and negative charged amino acids (yellow) in amphiphilic α-helixes of PLIN2 as predicted by Heliquest. The structural comparison was performed using PyMOL. (B) Residue E73 is in a different structural domain from the corresponding amino acid of PLIN3, which is in a coil. (C) Heliquest also predicted that E73 of PLIN2 is in an amphiphilic α-helix while E86 of PLIN3 is not. (D) The amino acid sequences of PLIN2 and PLIN3 were compared by ClustalX and shown by ESPript 3.0. The triangles represent negatively charged amino acids in the amphiphilic α-helixes of PLIN2. The red triangle shows the site of E73. The sequences marked by a-f were structurally different between PLIN2 and PLIN3.

According to previous research, the α-helix bundles play an important role in the targeting of PLIN2 to LDs (Najt et al., 2014). The structure reveals several sites of negatively charged amino acids in the amphiphilic α-helixes that may influence binding in the presence of the anionic PI. Protein interaction with PI is a complex phenomenon, involving multiple factors like electrostatic interaction, repulsive desolvation, and the hydrophobic-hydrophobic interaction between proteins and phospholipid layer. It is believed that electrostatic interaction is the most dominant it in this situation (Mulgrew-Nesbitt et al., 2006). The glutamic acid residue at 73 position (Figure 4A) in PLIN2 and the corresponding glutamic acid in PLIN3 at 86 position are of particular interest (Figure 4B and C). A close-up comparison is shown in Figure 4D where the amino acid sequences of PLIN2 and PLIN3 were compared by ClustalX and shown by ESPript 3.0 (Robert and Gouet, 2014). E73 of PLIN2 is located in the potential membrane targeting α-helix, whereas E83 of PLIN3 is in a coil in a spatial configuration away from the phospholipid surface (Rowe et al., 2016). Therefore, E73 may contribute to the targeting of PLIN2 but E83 cannot serve the same purpose in PLIN3.

To test this hypothesis, various mutants of PLIN2 were constructed where negatively charged amino acids were replaced with positively charged or neutral amino acids, *i*.*e*., asparagine, glutamine or lysine. These negatively charged amino acids were labeled in the α-helixes of PLIN2 sequence alignment (Figure 4D). The bacteria lysate expressing PLIN2 mutants was incubated with adiposomes. In Figure 5A, Western blot results of each PLIN2 mutant were compared to the wild type PLIN2. In Figure 5B, the chart summarizes the change ratio of each mutant compared to the control, quantified by gray scale intensity of the bands in the Western blot, using the following equation:

**Figure 5.**
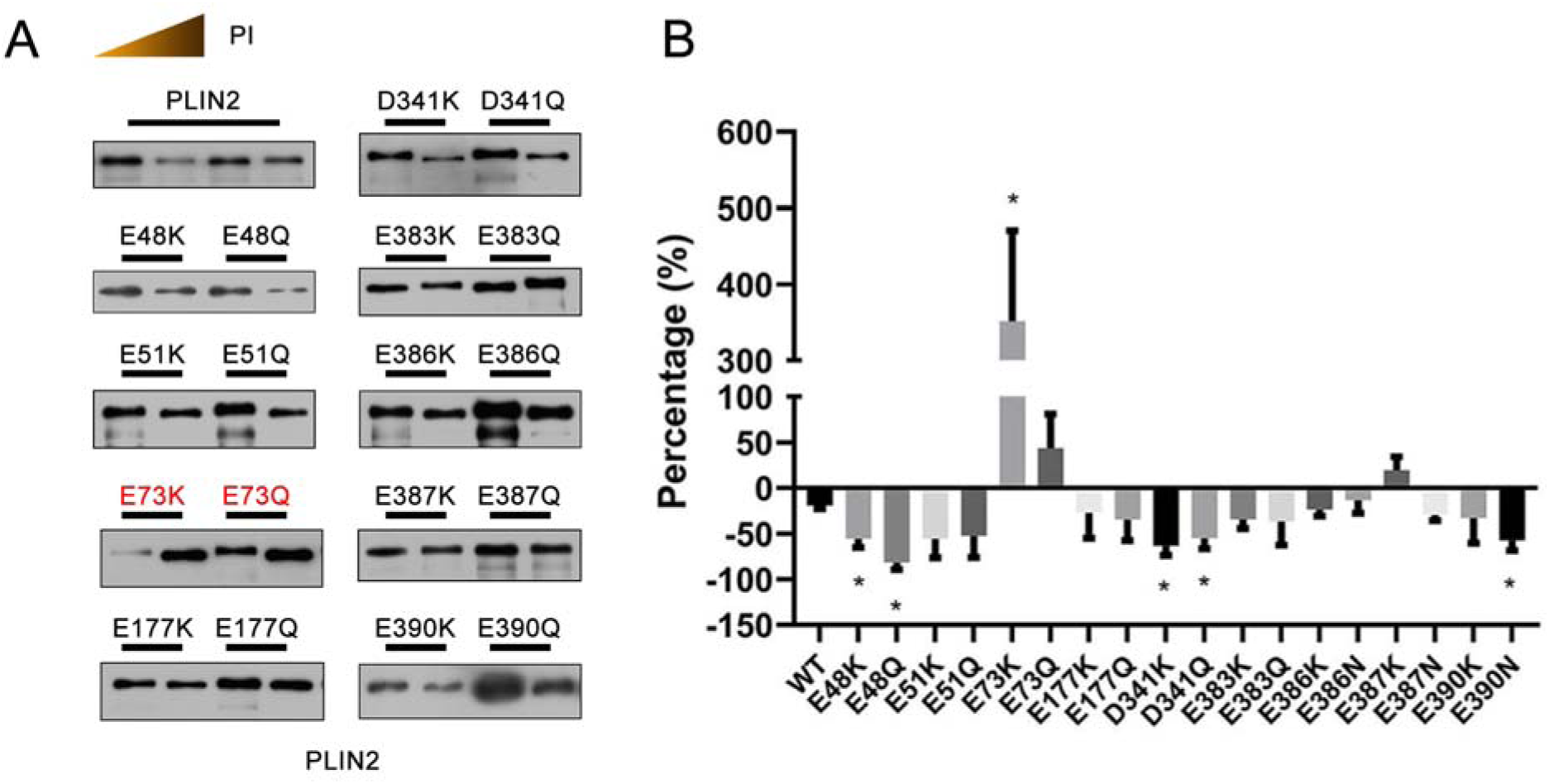
The 73^rd^ glutamic acid of PLIN2 affects its targeting to adiposomes in the presence of PI. Negatively charged amino acids in the amphiphilic α-helixes of PLIN2 were mutated to positively charged or neutral amino acids. All Western blots were performed using anti-PLIN2 antibody. The mutants were expressed in Transetta (DE3) following the method described in *Materials and Methods*. (A) After two washes of the adiposomes, proteins were analyzed by Western Blot. Only mutant E73K had increased targeting to adiposomes composed of 92.4% DOPC + 7.6% PI. (B) The densitometry from three independent experiments was analyzed by ImageJ. Data represent mean ± s.e.m., n = 3. \**P* < 0.05, two-tailed *t*-test.

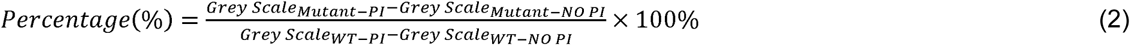

where *grey scale*_*Mutant-PI*_ is the intensity of mutant sample in the presence of PI, *grey scale*_*mutant-NO* *PI*_ is the intensity of mutant sample in the absence of PI, *grey scale*_*WT-PI*_ is the intensity of wild type sample in the presence of PI, and *grey scale*_*WT-NO* *PI*_ is the intensity of wild type sample in the absence of PI. Binding was significantly enhanced when the E73 was replaced by lysine. This result suggests that there is a potential role of for charge interactions in PLIN2 binding to PI containing monolayer membrane. The binding was slightly enhanced when E73 was mutated to glutamine, which also suggests that the binding at this site was affected by charge. The results confirm that E73 has an inhibitory role in PLIN2 binding on PI containing adiposome membranes.

The LD binding ability of the PLIN2 mutants was also tested in cells. Mutants at residues E48, E73, and D341 were overexpressed in Huh7 cells and all of them targeted LDs similar to wild type (data not shown). As shown in Figure S5 all of the mutants bound LDs without any noticeable change. Thus, none of these charged amino acids had an effect on the LD targeting ability of PLIN2 under intracellular conditions. Therefore, the inhibitory action against PI containing membranes may only regulate PLIN2 localization in situations where the concentration of PI changes and no other proteins are involved. Compared with PC and PE, PI is more negatively charged at physiological pH (≈7.4) and can carry multiple negative charges once phosphorylated (Marsh, 2013). The targeting of proteins to LDs is potentially influenced by the ionized phospholipid head group on the LD monolayer membrane.

### Pl is important for TAG hydrolase activity of ATGL on adiposomes

ATGL is a well-known TAG lipase that can translocate to LDs (Bartz et al., 2007b; Zimmermann et al., 2004). The main function of ATGL on LDs is to catalyze the first step of TAG hydrolysis (Zimmermann et al., 2004). Figure S6A shows that recombinant ATGL could target adiposomes without interacting with other proteins, indicating that ATGL recognizes the adiposome monolayer membrane. Active ATGL is abundant in the cytosol of brown adipose tissue (BAT) (Yu et al., 2015). BAT cytosol was incubated with adiposomes prepared using DOPC and PI or DOPC and DOPE (Figure S6B). There were no significant differences in ATGL targeting with increasing ratio of DOPE but targeting was slightly enhanced with an increasing ratio of PI. A diagram of an ATGL enzymatic activity assay is shown in Figure 6A. To study the effect of PI on the activity of ATGL *in vitro*, substrates were optimized using BAT cytosol as the enzyme source. High concentrations of imidazole (500 mM), removal of PI (only PC on the surface of adiposome), and Buffer B, all led to decreased enzymatic activity of ATGL in BAT cytosol (Figure 6B). The activity of the purified ATGL was very low (data not shown). Therefore, experiments were performed using bacteria lysate expressing ATGL, with BAT cytosol as a positive control. An increased ratio of PI promoted the enzymatic activity of ATGL from BAT cytosol (Figure 6C). The activity was significantly increased when the ratio of PI reached 25%. This result indicates that PI played a role in stimulating the activity of ATGL.

**Figure 6.**
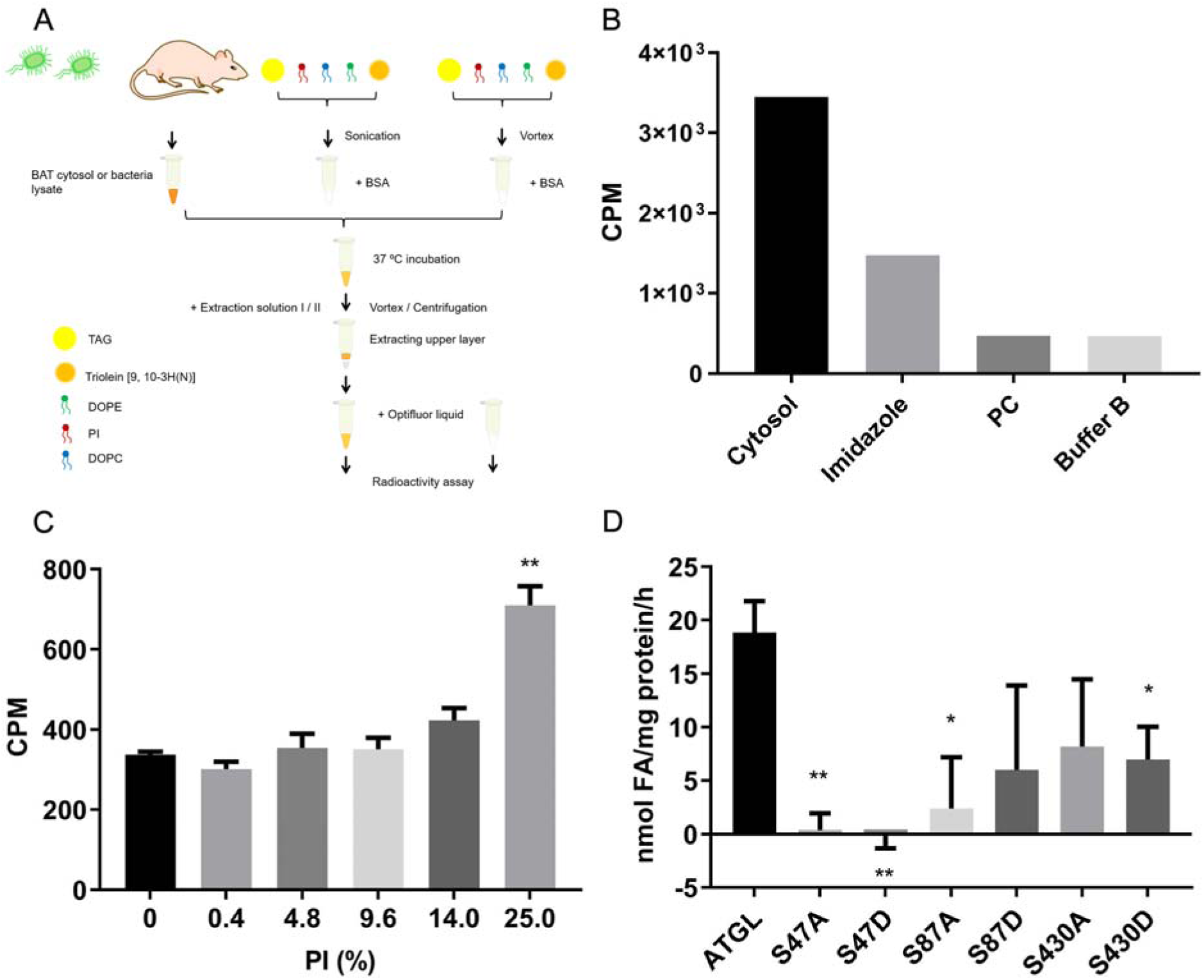
Enzymatic assay of ATGL using adiposomes. (A) Schematic of the experimental design. Triolein [9, 10-3H(N)] was used as a radio labeled substrate to perform enzyme activity assays. Equal amounts of BAT cytosol were incubated with sonication prepared emulsion containing increasing proportions of PI. (B) High concentrations of imidazole, removal of PI (adiposomes composed solely of PC), and inclusion of Buffer B led to decreased enzymatic activity. (C) The highest ATGL activity was obtained with the sonication prepared substrate composed of 25% PI. Data represent mean ± s.e.m., n = 3. \**P* < 0.05, two-tailed *t*-test. (D) Radio labeled adiposomes were prepared and incubated with bacteria lysate expressing ATGL mutants. The lipolytic activities of S47A, S47D, and S87A mutants were significantly decreased compared to the control. Data represent mean ± s.e.m., n = 3. \**P* < 0.05, \*\**P* < 0.01, two-tailed t-test.

The adiposome system offers a simple means to study the activity of ATGL. ATGL is regulated by phosphorylation with many phosphorylation sites having been identified (Ahmadian et al., 2011; Bartz et al., 2007b; Pagnon et al., 2012; Xie et al., 2014). ATGL mutants were constructed by site-directed mutation to mimic the phosphorylated (aspartic acid) or non-phosphorylated (alanine) protein. Compared with the wild type ATGL, the activity of mutants S47A, S47D, S87A, S87D, S430A and S430D were measured (Figure 6D). The activity of mutants decreased in all cases and decreased most significantly for mutants S47A, S47D and S87A. Next, ATGL was knocked out in C2C12 cells using the CRISPR-Cas9 system (Figure S7). Monoclone lines KO-2-15 were the cells transfected with a target sequence but did not show frameshift mutation (control), whereas the monoclone lines KO-2-16 was transfected with another target sequence and found missing one nucleotide and causing the frameshift mutation. The cells were fractionated and analyzed by silver stained SDS-PAGE and Western blot (Figure S7A, B and C). Both the protein profile and Western blot confirmed the knockout of ATGL in the KO-2-16 cells, as ATGL was highly expressed in KO-2-15 than KO-2-16. There was reduced PLIN2 in the knockout cells (Figure S7C) The mitochondria and LDs were larger in the knockout cells, demonstrating the important role of ATGL in neutral lipid degredation (Figure S7D).

The S47A, S47D, S87A, and S87D mutants were overexpressed in C2C12 cells followed by immunofluorescence detection using anti-Flag antibody. Fluorescent micrographs revealed that the S47A and the S47D mutants had reduced lipolytic activity compared to wild type ATGL, based on the size of LDs. Lipolytic activity of the S47D mutant was particularly suppressed (Figure 7). Interestingly, there was no difference in the apparent lipolytic capacity of the S87A mutant with or without oleate treatment while the activity of the S87D mutant increased with oleate treatment, but was still lower than wild type (data not shown). The activity of the S430A and S430D mutants were not tested by immunofluorescence. These results demonstrate that S47 is an important active site and phosphorylation might not be the only factor determining enzymatic activity. CGI-58 is known to target LDs after oleate treatment (Liu et al., 2004). Therefore, CGI-58 might bind to S87D instead of S87A to activate the TAG hydrolase activity.

**Figure 7.**
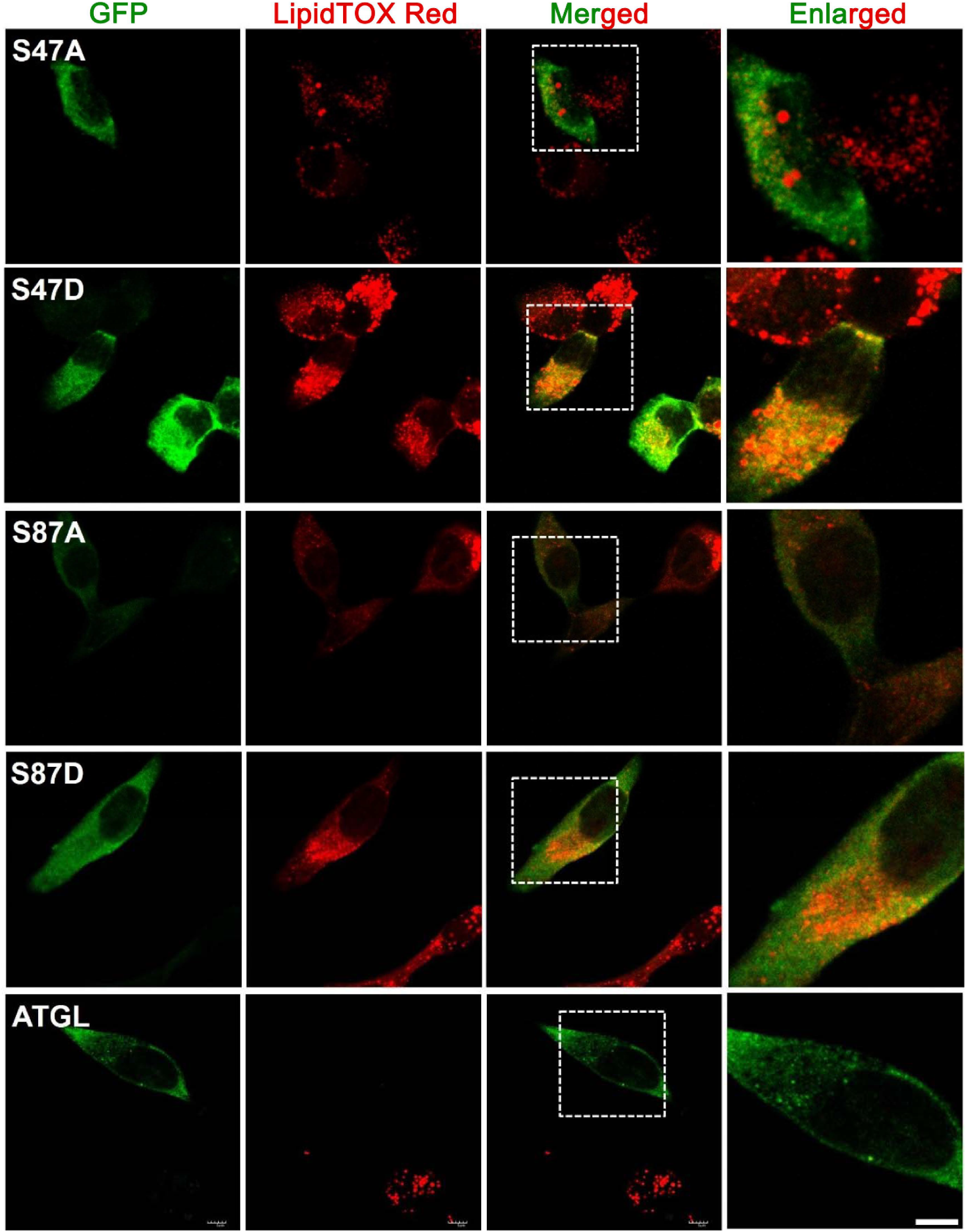
TAG hydrolase activity of ATGL is reduced in the mutants of S47 or S87. Wild type ATGL and mutants were overexpressed in C2C12 cells. ATGL was stained by immunofluorescence using mouse anti-Flag followed by anti-mouse FITC coupled antibody. Lipid droplets were stained with LipidTOX Red. Mutations of S47 or S87 caused decreased enzymatic activity, thus leaving more lipid droplets compared to wild type ATGL. Scale bar, 5 μm.

## DISCUSSION

The high purity, structural similarity and diameter homogeneity of adiposomes prepare them as the ideal model for mimicking LD *in vitro*. They therefore underlie the new methodology to characterize the interaction between LD and proteins. In this study it was assessed whether the differences in targeting of PI containing adiposomes by PLIN2 and PLIN3 originated from the negatively charged amino acids in the membrane binding α-helixes. The scatchard plots clearly show that the two LD associated proteins are able to saturate on the surface of adiposomes and the binding behavior can be quantitively analyzed using adiposome. This is the first time to report the characterization of LD specific protein binding affinity on LD *in vitro* model. Compared to previous researches on LD specific protein binding mechanism, it provides new sights for quantitative dynamics analysis, which is different from the qualitative analysis (Nakamura and Fujimoto, 2003; Rowe et al., 2016; Sletten et al., 2014). By analyzing the binding dynamics, PI was found to be a significant factor affecting the binding of PLIN2 on adiposomes. The behavior of PLIN2 mutants showed the importance of E73 in regulating targeting to adiposomes in the presence of PI, which may be related to charge repulsion. Under physiological pH, the phosphatidylcholine surface of adiposomes is a zwitterion that carries a negative charge on the phosphate moiety and positive charge on the choline group. The latter is less accessible due to its bulk, which may result in the negative charge dominating interactions with other charged species. Cations in the cellular milieu can penetrate membranes past choline headgroups to interact with phosphate and ester groups. PI contributes a greater density of negative charge as well as a bulky sugar ring containing five hydroxyl groups (Mak, 2013). Thus, the excess charge of PI could potentially repel E73 in the α-helix of PLIN2.

While the predicted structure suggests that E73 is in a position to interact with the LD membrane, it is not in a domain that has been implicated in membrane targeting. Truncation studies showed that the amino-terminal sequence (aa 1-181) and the carboxyl-terminal sequence (aa 277-426) are pivotal for PLIN2 to target LDs (Nakamura and Fujimoto, 2003). The conserved PAT domain, a more than 100 amino acids region close to the N-termini of PAT family proteins, is unnecessary for PLIN2 targeting (Garcia et al., 2003; McManaman et al., 2003; Najt et al., 2014). E73 localizes in the PAT domain of PLIN2, and not the domains identified as directly responsible for LD binding. Thus, it is an unexpected outcome in this study that this amino acid influences PLIN2 targeting to adiposomes. This result may derive from the balance of nonspecific electrostatic, desolvation, and non-polar interactions between the protein and phospholipid layer (Mulgrew-Nesbitt et al., 2006). The latter two effects act on a short range while the electrostatic interaction works on long range. Hence, glutamic acid residues localized in the targeting domains of PLIN2 are more affected by desolvation and hydrophobic interaction since these hydrophobic helixes are likely embedded in the phospholipid acyl-chain region. This could potentially affect the protonation (charged) state as the buffering effect of the physiological solution is lost. (Kory et al., 2016; Mulgrew-Nesbitt et al., 2006). In contrast, the E73 of the PAT domain of PLIN2 is more exposed and thus retains charge. Therefore, it would be more affected by Coulombic force, such that the mutants show significant changes in targeting when the charge-charge repulsion is altered.

For PE, the conical molecule introduces a negative curvature to the lipid surface (van den Brink-van der Laan et al., 2004; Zanghellini et al., 2010). Compared to PC, the projections of PE are up to 15±7 Å^2^ larger than PC, which leads to more packing defects. This is conducive to the insertion of peripheral membrane proteins into the lipid membrane surface (van den Brink-van der Laan et al., 2001). For example, the binding of apoLp-III, an amphipathic α-helix bundle protein, to a phospholipid-coated oil surface is promoted by increasing concentrations of PE in a PC/PE mixture (Mirheydari et al., 2018). Thus, lipid density and packing effects offer a straightforward explanation for the increased PLIN2 binding to PC/PE adiposomes. The saturation of PLINs may also correspond to the packing defects on adiposomes. Potentially, the packing defects occupying by PLIN2 are more than same concentration of PLIN3. Interestingly, ATGL maintains a constant binding density on adiposomes of varied PE content (Figure S6B), suggesting a different binding mechanism from PLIN2. In sum, the binding of PLIN2 is regulated by the conjugation of amphipathic helix and lipid packing defects, and the electrostatic interaction between charged residues and charged phospholipids on the surface of adiposome.

To study the lipase activity of ATGL, cytosolic extracts containing the enzyme were incubated with either the purified LD, LD mimics (*i*.*e*. a neutral lipid and buffer mixture), or neutral lipid and phospholipid microdroplets (Duncan et al., 2008; Schweiger et al., 2008; Zimmermann et al., 2004). However, these methods have limitations in that the composition of isolated LDs cannot be manipulated easily and the surfaces of the synthetic substrates are dissimilar from natural LDs. In contrast, the adiposome creates a surface which is close to the structure of LDs with a homogeneous size distribution such that it may more closely reflect the native lipase activity (Figure S2F). In this study the results obtained with ATGL mutants suggest that the phosphorylation state of ATGL affects the *in vitro* lipolysis capacity. Serine 47 has been reported as a phosphorylation site necessary for TAG hydrolysis (Duncan et al., 2010; Lake et al., 2005). Our results support this finding since the hydrolase activity of the serine 47 mutants was decreased; the fluorescence images show no decrease in the size of LDs. Serine 430 was found to be neither critical for LD targeting nor necessary for TAG hydrolysis (Duncan et al., 2010). However, our data indicates that serine 430 can decrease the *in vitro* lipase activity of ATGL. The S87A mutant also showed a decline in lipase activity which few publications have mentioned.

In previous work the preparation of adiposomes was described in detail, and the structure and LD protein binding ability of adiposomes was verified (Wang et al., 2016). This work delivers further evidence that adiposomes provide a suitable platform for *in vitro* study of LD proteins. Compared to the emulsion prepared by homogenizing phospholipids and neutral lipids, purified adiposomes show a substantial advantage by mimicking the actual structure of LDs. In summary, adiposomes were used as an *in vitro* model of LDs for the study of LD-associated proteins. The proteins studied, PLIN2, PLIN3, and ATGL, bound the adiposomes. The addition of PI to the phospholipid composition decreased PLIN2 targeting, but not PLIN3 or ATGL. This suggests that different mechanisms are responsible for the targeting of these proteins. PLIN2 and PLIN3 binding was saturable and the binding properties were analyzed with Scatchard plots. A comparison of the structures of PLIN2 and PLIN3 suggests that the E73 residue in the PAT domain of PLIN2 influences the targeting of PI containing membranes. In contrast, the analogous E86 residue in PLIN3 is located in a β-turn, which may explain the different response to PI content. ATGL displayed increasing lipolytic activity against PI containing adiposomes. Phosphorylation site mutants in ATGL were examined and serine 47 was found to be a significant enzyme active site and phosphorylation of serine 87 was required to maintain activity. This work showcases the utility of using adiposomes as an *in vitro* LD model to study the targeting of LD-associated proteins and the determination of lipase activity.

### Limitations of the Study

In this present study, we apply adiposome as the LD *in vitro* model, to develop a method determining the LD specific protein binding affinity. We also prove the availability of adiposome on determining the *in vitro* lipase activity of ATGL. However, it is a proof of concept recently so a vast amount of proteins may be needed to further support this study. Likewise, we do not include more lipid components of adiposome than PI and PE in present study.

## METHODS

All methods can be found in the accompanying Transparent Methods supplemental file.

## ABBREVIATIONS

DOPC: 1,2-dioleoyl-sn-glycero-3-phosphocholine.
DOPE: 1,2-dioleoyl-sn-glycero-3-phosphoethanolamine.
PI: phosphatidylinositol.
Liver PI: L-α-phosphatidylinositol (Liver, Bovine) (sodium salt).
TAG: triacylglycerol.
ADRP: adipose differentiation-related protein.
TIP47: tail-interacting protein of 47 kDa.
PLINs: perilipin family proteins.
ATGL: adipose triglyceride lipase.
ALD: artificial lipid droplet.
LD: lipid droplet.
FFF-MALS: field flow fractionation-multi-angle light scattering.
TEM: transmission electron microscope.

## SUPPLEMENTAL INFORMATION

Supplemental Information can be found online.

## Deposited Data

The original data of this study is included in https://data.mendeley.com/datasets/yyvxtx3bmc/draft?a=f8310a45-8ccb-4a0b-9674-a9b77a7a1941

## ACKNOWLEDGEMENTS

The authors thank Dr. John Zehmer for his critical reading and useful suggestions, Dr. Yang Wang for advices on adiposome preparation, and Dr. Liujuan Cui for administrative and technique support. This work was supported by the National Key R&D Program of China (Grant No. 2016YFA0500100, 2018YFA0800700 and 2018YFA0800900), National Natural Science Foundation of China (Grant No. 91857201, 91954108, 31671402, 31671233, 31701018 and U1702288). This work was also supported by the “Personalized Medicines—Molecular Signature-based Drug Discovery and Development”, Strategic Priority Research Program of the Chinese Academy of Sciences, Grant No. XDA12040218.

## AUTHOR CONTRIBUTIONS

A.M. and P.L. conceived the study and designed the experiments. X.M., Z.Z, S.Z. and C.Z. performed the experiments and analyzed the data. P.L. analyzed data and provided guidance and support. X.M., Z.Z, A.M. and P.L. wrote the paper.

## DECLARATION OF INTERESTS

The authors declare that there is no conflict of interests regarding the publication of this article.

## Notes

https://data.mendeley.com/datasets/yyvxtx3bmc/draft?a=f8310a45-8ccb-4a0b-9674-a9b77a7a1941

### Competing Interest Statement

The authors have declared no competing interest.

### Summary of Updates

New strategies to decode the specific protein targeting mechanism on lipid droplet (LD) are urgently needed. Using adiposome, the LD binding of perilipin 2 (PLIN2), perilipin 3 (PLIN3), and adipose triglyceride lipase (ATGL) were studied. Scatchard analysis found that the binding of PLIN2 to the adiposome surface was saturable, pointing to a specific membrane binding partner. Phosphatidylinositol (PI) was found to inhibit PLIN2 binding while it did not impede PLIN3. Structural analysis combined with mutagenesis revealed that the 73rd glutamic acid of PLIN2 is significant for the effect of PI on the protein binding. The presence of PI significantly stimulated the activity of ATGL in vitro. The phosphorylation site mutants of ATGL were found reducing the lipase activity in the adiposome system. Our study demonstrates the utility of adiposome as a powerful, manipulatable model system for the characterization of LD binding and enzymatic activity of LD proteins in vitro.

## REFERENCES

Ahmadian, M., Abbott, Marcia J., Tang, T., Hudak, Carolyn S.S., Kim, Y., Bruss, M., Hellerstein, Marc K., Lee, H.-Y., Samuel, Varman T., Shulman, Gerald I., et al. (2011). Desnutrin/ATGL Is regulated by AMPK and is required for a brown adipose phenotype. Cell Metab 13, 739–748.

Barneda, D., Planas-Iglesias, J., Gaspar, M.L., Mohammadyani, D., Prasannan, S., Dormann, D., Han, G.-S., Jesch, S.A., Carman, G.M., Kagan, V., et al. (2015). The brown adipocyte protein CIDEA promotes lipid droplet fusion via a phosphatidic acid-binding amphipathic helix. eLife 4, e07485.

Bartz, R., Li, W.-H., Venables, B., Zehmer, J.K., Roth, M.R., Welti, R., Anderson, R.G.W., Liu, P., and Chapman, K.D. (2007a). Lipidomics reveals that adiposomes store ether lipids and mediate phospholipid traffic. J Lipid Res 48, 837–847.

Bartz, R., Zehmer, J.K., Zhu, M., Chen, Y., Serrero, G., Zhao, Y., and Liu, P. (2007b). Dynamic activity of lipid droplets: protein phosphorylation and GTP-mediated protein translocation. J Proteome Res 6, 3256–3265.

Bersuker, K., and Olzmann, J.A. (2017). Establishing the lipid droplet proteome: Mechanisms of lipid droplet protein targeting and degradation. Biochimica et biophysica acta Molecular and cell biology of lipids 1862, 1166–1177.

Boeszoermenyi, A., Nagy, H.M., Arthanari, H., Pillip, C.J., Lindermuth, H., Luna, R.E., Wagner, G., Zechner, R., Zangger, K., and Oberer, M. (2015). Structure of a CGI-58 motif provides the molecular basis of lipid droplet anchoring. J Biol Chem 290, 26361–26372.

Bulankina, A.V., Deggerich, A., Wenzel, D., Mutenda, K., Wittmann, J.G., Rudolph, M.G., Burger, K.N., and Höning, S. (2009). TIP47 functions in the biogenesis of lipid droplets. J Cell Biol 185, 641–655.

Chen, Y., Jena, K.C., Lütgebaucks, C., Okur, H.I., and Roke, S. (2015). Three Dimensional Nano “Langmuir Trough” for Lipid Studies. Nano Letters 15, 5558–5563.

Chong, B.M., Russell, T.D., Schaack, J., Orlicky, D.J., Reigan, P., Ladinsky, M., and McManaman, J.L. (2011). The adipophilin C terminus is a self-folding membrane-binding domain that is important for milk lipid secretion. J Biol Chem 286, 23254–23265.

Ding, Y., Yang, L., Zhang, S., Wang, Y., Du, Y., Pu, J., Peng, G., Chen, Y., Zhang, H., Yu, J., et al. (2012). Identification of the major functional proteins of prokaryotic lipid droplets. J Lipid Res 53, 399–411.

Duncan, R.E., Sarkadi-Nagy, E., Jaworski, K., Ahmadian, M., and Sul, H.S. (2008). Identification and functional characterization of adipose-specific phospholipase A2 (AdPLA). J Biol Chem 283, 25428–25436.

Duncan, R.E., Wang, Y., Ahmadian, M., Lu, J., Sarkadi-Nagy, E., and Sul, H.S. (2010). Characterization of desnutrin functional domains: critical residues for triacylglycerol hydrolysis in cultured cells. J Lipid Res 51, 309–317.

Farese, R.V., and Walther, T.C. (2009). Lipid droplets finally get a little R-E-S-P-E-C-T. Cell 139, 855–860.

Fei, W., Shui, G., Zhang, Y., Krahmer, N., Ferguson, C., Kapterian, T.S., Lin, R.C., Dawes, I.W., Brown, A.J., Li, P., et al. (2011). A role for phosphatidic acid in the formation of “supersized” lipid droplets. PLoS Genet 7, e1002201.

Garcia, A., Sekowski, A., Subramanian, V., and Brasaemle, D.L. (2003). The central domain is required to target and anchor perilipin A to lipid droplets. J Biol Chem 278, 625–635.

Hickenbottom, S.J., Kimmel, A.R., Londos, C., and Hurley, J.H. (2004). Structure of a lipid droplet protein: the PAT family member TIP47. Structure 12, 1199–1207.

Huang, C.Y., and Huang, A.H.C. (2017). Unique Motifs and Length of Hairpin in Oleosin Target the Cytosolic Side of Endoplasmic Reticulum and Budding Lipid Droplet. Plant Physiol 174, 2248–2260.

Kory, N., Farese, R.V., Jr., and Walther, T.C. (2016). Targeting Fat: Mechanisms of Protein Localization to Lipid Droplets. Trends Cell Biol 26, 535–546.

Krahmer, N., Guo, Y., Wilfling, F., Hilger, M., Lingrell, S., Heger, K., Newman, H.W., Schmidt-Supprian, M., Vance, D.E., Mann, M., et al. (2011). Phosphatidylcholine synthesis for lipid droplet expansion is mediated by localized activation of CTP:phosphocholine cytidylyltransferase. Cell Metab 14, 504–515.

Lake, A.C., Sun, Y., Li, J.-L., Kim, J.E., Johnson, J.W., Li, D., Revett, T., Shih, H.H., Liu, W., Paulsen, J.E., et al. (2005). Expression, regulation, and triglyceride hydrolase activity of Adiponutrin family members. J Lipid Res 46, 2477–2487.

Lemmon, M.A. (2008). Membrane recognition by phospholipid-binding domains. Nat Rev Mol Cell Biol 9, 99–111.

Leung, K.F., Baron, R., Ali, B.R., Magee, A.I., and Seabra, M.C. (2007). Rab GTPases containing a CAAX motif are processed post-geranylgeranylation by proteolysis and methylation. J Biol Chem 282, 1487–1497.

Liu, P., Ying, Y., Zhao, Y., Mundy, D.I., Zhu, M., and Anderson, R.G.W. (2004). Chinese hamster ovary K2 cell lipid droplets appear to be metabolic organelles involved in membrane traffic. J Biol Chem 279, 3787–3792.

Liu, W., Xie, Y., Ma, J., Luo, X., Nie, P., Zuo, Z., Lahrmann, U., Zhao, Q., Zheng, Y., Zhao, Y., et al. (2015). IBS: an illustrator for the presentation and visualization of biological sequences. Bioinformatics 31, 3359–3361.

Mak, L.H. (2013). Lipid Signaling and Phosphatidylinositols. In Encyclopedia of Biophysics, G.C.K. Roberts, ed. (Berlin, Heidelberg: Springer Berlin Heidelberg), pp. 1286–1289.

Marsh, D. (2013). Handbook of lipid bilayers (CRC press).

Martin, S., and Parton, R.G. (2006). Lipid droplets: a unified view of a dynamic organelle. Nat Rev Mol Cell Biol 7, 373–378.

McManaman, J.L., Zabaronick, W., Schaack, J., and Orlicky, D.J. (2003). Lipid droplet targeting domains of adipophilin. J Lipid Res 44, 668–673.

Mirheydari, M., Mann, E.K., and Kooijman, E.E. (2018). Interaction of a model apolipoprotein, apoLp-III, with an oil-phospholipid interface. Biochim Biophys Acta Biomembr 1860, 396–406.

Miura, S., Gan, J.-W., Brzostowski, J., Parisi, M.J., Schultz, C.J., Londos, C., Oliver, B., and Kimmel, A.R. (2002). Functional conservation for lipid storage droplet association among perilipin, ADRP, and TIP47 (PAT)-related proteins in mammals, drosophila and dictyostelium. J Biol Chem 277, 32253–32257.

Mulgrew-Nesbitt, A., Diraviyam, K., Wang, J., Singh, S., Murray, P., Li, Z., Rogers, L., Mirkovic, N., and Murray, D. (2006). The role of electrostatics in protein–membrane interactions. Biochimica et biophysica acta Molecular and cell biology of lipids 1761, 812–826.

Na, H., Zhang, P., Chen, Y., Zhu, X., Liu, Y., Liu, Y., Xie, K., Xu, N., Yang, F., Yu, Y., et al. (2015). Identification of lipid droplet structure-like/resident proteins in Caenorhabditis elegans. Biochim Biophys Acta Mol Cell Res 1853, 2481–2491.

Najt, C.P., Lwande, J.S., McIntosh, A.L., Senthivinayagam, S., Gupta, S., Kuhn, L.A., and Atshaves, B.P. (2014). Structural and functional assessment of perilipin 2 lipid binding domain(s). Biochemistry 53, 7051–7066.

Nakamura, N., and Fujimoto, T. (2003). Adipose differentiation-related protein has two independent domains for targeting to lipid droplets. Biochem Bioph Res Co 306, 333–338.

Ohsaki, Y., Suzuki, M., and Fujimoto, T. (2014). Open questions in lipid droplet biology. Chemistry & biology 21, 86–96.

Pagnon, J., Matzaris, M., Stark, R., Meex, R.C.R., Macaulay, S.L., Brown, W., O’Brien, P.E., Tiganis, T., and Watt, M.J. (2012). Identification and functional characterization of protein kinase A phosphorylation sites in the major lipolytic protein, adipose triglyceride lipase. Endocrinology 153, 4278–4289.

Phan, T.K., Lay, F.T., Poon, I.K.H., Hinds, M.G., Kvansakul, M., and Hulett, M.D. (2016). Human β-defensin 3 contains an oncolytic motif that binds PI(4,5)P2 to mediate tumour cell permeabilisation. Oncotarget 7, 2054–2069.

Poppelreuther, M., Sander, S., Minden, F., Dietz, M.S., Exner, T., Du, C., Zhang, I., Ehehalt, F., Knüppel, L., Domschke, S., et al. (2018). The metabolic capacity of lipid droplet localized acyl-CoA synthetase 3 is not sufficient to support local triglyceride synthesis independent of the endoplasmic reticulum in A431 cells. Biochimica et biophysica acta Molecular and cell biology of lipids 1863, 614–624.

Prévost, C., Sharp, M.E., Kory, N., Lin, Q., Voth, G.A., Farese, R.V., Jr., and Walther, T.C. (2018). Mechanism and determinants of amphipathic helix-containing protein targeting to lipid droplets. Dev Cell 44, 73-86.e74.

Robert, X., and Gouet, P. (2014). Deciphering key features in protein structures with the new ENDscript server. Nucleic Acids Res 42, W320–W324.

Rowe, E.R., Mimmack, M.L., Barbosa, A.D., Haider, A., Isaac, I., Ouberai, M.M., Thiam, A.R., Patel, S., Saudek, V., Siniossoglou, S., et al. (2016). Conserved Amphipathic Helices Mediate Lipid Droplet Targeting of Perilipins 1-3. J Biol Chem 291, 6664–6678.

Schweiger, M., Schoiswohl, G., Lass, A., Radner, F.P.W., Haemmerle, G., Malli, R., Graier, W., Cornaciu, I., Oberer, M., Salvayre, R., et al. (2008). The C-terminal region of human adipose triglyceride lipase affects enzyme activity and lipid droplet binding. J Biol Chem 283, 17211–17220.

Shen, W.J., Patel, S., Miyoshi, H., Greenberg, A.S., and Kraemer, F.B. (2009). Functional interaction of hormone-sensitive lipase and perilipin in lipolysis. J Lipid Res 50, 2306–2313.

Sletten, A., Seline, A., Rudd, A., Logsdon, M., and Listenberger, L.L. (2014). Surface features of the lipid droplet mediate perilipin 2 localization. Biochem Bioph Res Co 452, 422–427.

Subramanian, V., Garcia, A., Sekowski, A., and Brasaemle, D.L. (2004). Hydrophobic sequences target and anchor perilipin A to lipid droplets. J Lipid Res 45, 1983–1991.

Tauchi-Sato, K., Ozeki, S., Houjou, T., Taguchi, R., and Fujimoto, T. (2002). The surface of lipid droplets is a phospholipid monolayer with a unique fatty acid composition. J Biol Chem 277, 44507–44512.

Tzen, J.T., and Huang, A.H. (1992). Surface structure and properties of plant seed oil bodies. J Cell Biol 117, 327.

van den Brink-van der Laan, E., Antoinette Killian, J., and de Kruijff, B. (2004). Nonbilayer lipids affect peripheral and integral membrane proteins via changes in the lateral pressure profile. Biochim Biophys Acta Biomembr 1666, 275–288.

van den Brink-van der Laan, E., Dalbey, R.E., Demel, R.A., Killian, J.A., and de Kruijff, B. (2001). Effect of nonbilayer lipids on membrane binding and insertion of the catalytic domain of leader peptidase. Biochemistry 40, 9677–9684.

Walther, T.C., Chung, J., and Jr., R.V.F. (2017). Lipid droplet biogenesis. Curr Opin Cell Biol 33, 491–510.

Walther, T.C., and Farese, R.V., Jr. (2012). Lipid droplets and cellular lipid metabolism. Annu Rev Biochem 81, 687–714.

Wang, Y., Zhou, X.-M., Ma, X., Du, Y., Zheng, L., and Liu, P. (2016). Construction of nanodroplet/adiposome and artificial lipid droplets. ACS Nano 10, 3312–3322.

Wolins, N.E., Brasaemle, D.L., and Bickel, P.E. (2006). A proposed model of fat packaging by exchangeable lipid droplet proteins. FEBS Lett 580, 5484–5491.

Xie, X., Langlais, P., Zhang, X., Heckmann, B.L., Saarinen, A.M., Mandarino, L.J., and Liu, J. (2014). Identification of a novel phosphorylation site in adipose triglyceride lipase as a regulator of lipid droplet localization. Am J Physiol Endocrinol Metab 306, E1449–E1459.

Xu, S., Zou, F., Diao, Z., Zhang, S., Deng, Y., Zhu, X., Cui, L., Yu, J., Zhang, Z., Bamigbade, A.T., et al. (2019). Perilipin 2 and lipid droplets provide reciprocal stabilization. Biophysics Reports 5, 145–160.

Yan, R., Qian, H., Lukmantara, I., Gao, M., Du, X., Yan, N., and Yang, H. (2018). Human SEIPIN binds anionic phospholipids. Dev Cell 47, 248-256.e244.

Yao, Y., Ding, L., and Huang, X. (2019). Diverse Functions of Lipids and Lipid Metabolism in Development. Small Methods, 1900564.

Yu, J., Zhang, S., Cui, L., Wang, W., Na, H., Zhu, X., Li, L., Xu, G., Yang, F., Christian, M., et al. (2015). Lipid droplet remodeling and interaction with mitochondria in mouse brown adipose tissue during cold treatment. Biochim Biophys Acta Mol Cell Res 1853, 918–928.

Zanghellini, J., Wodlei, F., and von Grünberg, H.H. (2010). Phospholipid demixing and the birth of a lipid droplet. J Theor Biol 264, 952–961.

Zhang, C., and Liu, P. (2017). The lipid droplet: A conserved cellular organelle. Protein & cell 8, 796–800.

Zhang, C., and Liu, P. (2019). The new face of the lipid droplet: lipid droplet proteins. Proteomics 19, e1700223.

Zhang, C., Yang, L., Ding, Y., Wang, Y., Lan, L., Ma, Q., Chi, X., Wei, P., Zhao, Y., Steinbüchel, A., et al. (2017). Bacterial lipid droplets bind to DNA via an intermediary protein that enhances survival under stress. Nat Commun 8, 15979.

Zimmermann, R., Strauss, J.G., Haemmerle, G., Schoiswohl, G., Birner-Gruenberger, R., Riederer, M., Lass, A., Neuberger, G., Eisenhaber, F., Hermetter, A., et al. (2004). Fat mobilization in adipose tissue is promoted by adipose triglyceride lipase. Science 306, 1383–1386.

