## Supplemental texts and figures for "Adiposome Targeting and Enzymatic Activity of Lipid Droplet-Specific Proteins"

### TRANSPARENT METHODS

#### Reagents

The following reagents were purchased from the indicated corporations: 1,2-dioleoyl-sn-glycero-3-phosphocholine (DOPC, 850375, Avanti), 1,2-dioleoyl-sn-glycero-3-phosphoethanolamine (DOPE, 850725, Avanti), L-α-phosphatidylinositol (Liver, Bovine) (sodium salt) (Liver PI, 840042, Avanti), LipidTOX Red Neutral Lipid Stain (H34476, Thermo Fisher), LipidTOX Green Neutral Lipid Stain (H34475, Thermo Fisher), MitoTracker Red CMXRos (M7512, Thermo Fisher), Hoechst 33258 (H21491, Thermo Fisher), Triolein [9, 10-3H(N)] (70805-83-3, PerkinElmer), Phenylmethylsulfonyl fluoride (PMSF, P7626, Millipore-Sigma), Colloidal Blue staining kit (LC6025, Invitrogen), Puromycin Dihydrochloride (A1113803, Invitrogen), Phusion® High-Fidelity DNA Polymerase (M0530S, New England Biolabs), BCA Protein Assay Kit (PI23227, Thermo Fisher), Anti-PLIN2 antibody (ab108323, Abcam), Anti-PLIN3 antibody (ab47638, Abcam), Anti-ATGL antibody (2138s, Cell Signaling), Anti-CGI-58 antibody (A6801, ABclonal). Triacylglycerol was acquired from rat fat pad, and the C57BL/6 mouse (male) were obtained from Vital River Laboratories. All animal protocols were approved by the Animal Care and Use Committee of the Institute of Biophysics and University of Chinese Academy of Sciences under the permission number SYXK（Jing）2016-0026. 25% glutaraldehyde solution (EM grade), Embed 812 kit, uranyl acetate and lead citrate were all purchased from Electron Microscopy Sciences (Hatfield, USA). Osmium tetraoxide (EM grade) was acquired from Nakalai Tesque.

#### Plasmids and primers

The primers used in this study can be found in Table S1. The plasmids can be found in Table S2.

#### Cell culture

Mouse C2C12 myoblasts and Huh7 cells were purchased from American Type Culture Collections and Shanghai Institutes for Biological Sciences, respectively. C2C12 myoblasts and human Huh7 hepatocarcinoma cells were maintained in DMEM supplemented with 10% (v/v) fetal bovine serum (FBS), 100 U $ml^{-1}$ penicillin, and 100 U $ml^{-1}$ streptomycin at 37°C with 5% $CO_2$.

**Experimental strains and subject details**

Transetta (DE3) strain was used for protein expression. TOP10 strain was used to construct recombinant DNA. Archives of the strains were maintained at -80ºC in 25% glycerol. The cell lines were cultured in the culture medium which consisted of high glucose Dulbecco's-Modified Eagle Medium (DMEM, C11965500BT, Invitrogen) containing 2 mM L-glutamine, 100 U $ml^{-1}$ penicillin, 100 μg $ml^{-1}$ streptomycin and 10% heat-inactivated fetal bovine serum (Gibco). Cultures were grown at 37ºC under 5% $CO_2$.

**Preparation of adiposomes and lipid emulsions**

The preparation of adiposomes followed the method reported previously (Wang et al., 2016). Two mg of total phospholipids (DOPC, DOPE, Liver PI) in chloroform were added to a 1.5 ml microcentrifuge tube, and the solvent was dried under a stream of nitrogen. Then 100 μl of Buffer B (20 mM HEPES, 100 mM KCl, 2 mM $MgCl_2$, pH 7.4) was added followed by 5 μl of TAG (extracted from rat fat pad in the laboratory). The tube containing lipids and buffer was vortexed for 24 cycles of 10 seconds on and 10 seconds off. Adiposomes in the milky emulsion were then purified using centrifugation as described previously. The purified adiposomes were analyzed by dynamic light scattering (DLS; Beckman Coulter). The concentration of adiposomes was measured by optical density at 600 nm (OD600) using an Eppendorf Biophotometer (Eppendorf) and EnSpire Multimode Plate Reader (Perkin Elmer). Particle counting was performed using the FFF-MALS system (Wyatt Technology). The lipid emulsion was prepared using the same materials while the mixture was sonicated using water bath sonicator for 6 min (1 min on and 10 seconds off).

**Ultrastructural analysis of adiposomes and emulsions by transmission electron microscopy (TEM)**

Briefly, adiposomes or emulsions were fixed with an equal volume of 2% glutaraldehyde in 0.1 M PB (pH 7.4) for 0.5 h at room temperature. Next an equal volume of 2% osmium tetraoxide was added to further fix the sample for 0.5 h at room temperature. The fixed adiposomes or emulsions were collected by centrifugation and processed for dehydration in an ascending concentration series of ethanol and infiltration subsequently in Embed 812. Afterwards, 70 nm sections were prepared with a Leica EM UC6 Ultramicrotome. The sections were then stained with uranyl acetate and lead citrate. The samples were observed with Tecnai Spirit electron microscope (FEI, Netherlands).

**Expression and purification of proteins**

A 6×-His tag was inserted at the N-terminus of a SMT3-PLIN2-GFP fusion protein expression vector which was constructed as previously reported (Wang et al., 2016). Standard molecular cloning techniques were used to fuse genes of PLIN3 and APPLE. PLIN3-APPLE was cloned into the pET28a expression vector and was expressed with an N-terminal 6×-His tag. Mutagenesis was conducted on the SMT3-PLIN2-GFP and SMT3-ATGL fusion protein expression vectors. All proteins were expressed in Transetta (DE3) in 2 × Yeast extract-Tryptone media. The expression and purification of proteins followed the published methods (Wang et al., 2016).

**Protein binding to adiposomes**

The previously published protein binding methods were slightly modified depending on the requirements of the different assays (Wang et al., 2016). Briefly, defined quantities of purified proteins were added to adiposome preparations in a final volume of 50 µl except in Scatchard analysis assays, which used a 60 µl volume. The mixture was gently vortexed three times, centrifuged at 1,000 × *g* for 10 seconds, and was then incubated at 37ºC in a water bath for 5 min or at 4ºC for 12 h for the Scatchard assay. The adiposome suspension was centrifuged at 21,130 × *g* for 5 min and the solution underneath was removed, while the top adiposome layer was reserved. The remaining adiposomes were then resuspended in 30 µl of Buffer B and centrifuged again. The wash procedure was repeated three times to remove any free proteins. The washed adiposomes were then used for microscopy or protein analysis.

**Fluorescence microscopy**

Adiposomes were incubated with LipidTOX Red (H34476, Invitrogen, 1:1,000 dilution, this ratio was fixed unless specifically mentioned) for 30 min at room temperature and then mounted on a slide. For protein containing samples, 5 µg SMT3-PLIN2-GFP was incubated with 30 µl adiposomes at 37°C for 5 min followed by three washes with 30 µl Buffer B with centrifugation at 21,130 × *g* to recover the adiposomes. Fluorescence images were obtained using an Olympus FV1000 confocal microscope, or a DeltaVision OMX V3 super resolution microscope.

For the immunofluorescence microscopy, C2C12 ATGL-Flag cells were seeded and grown overnight on a glass-bottomed plate. Then cells were placed on ice and washed three times with ice-cold PBS for 5 min each (140 mM NaCl, 2.7 mM KCl, 10 mM $Na_2HPO_4$, 1.8 mM $KH_2PO_4$, pH 7.4). Each additional step described below was followed by three washes. The cells were fixed in 4% paraformaldehyde at room temperature for 30 min and then permeabilized with 0.01% TritonX-100 in PBS at room temperature for 30 min. After blocking with 1% BSA in PBS at room temperature for 1 h, the cells were incubated with anti-Flag monoclonal antibody (1:100 diluted in 0.25% BSA/PBS) at room temperature for 1 h. Afterwards, the cells were washed using PBS and incubated with fluorescein isothiocyanat (FITC)-conjugated goat anti-rabbit IgG (1:100 diluted in 0.25% BSA/PBS) for 1 h at room temperature. LDs were stained using LipidTOX Red for 30 min. The coverslips were applied to a slide, mixed with 2 µl of mounting media, and sealed with nail polish. Cells were examined using an Olympus FV1000 confocal fluorescence microscope.

**Protein binding assay**

Adiposomes prepared in individual tubes were mixed together by gentle vortexing and were adjusted to OD600 = 20 using an Eppendorf Biophotometer. An equal volume of adiposomes (generally 30 µl) was added to 1.5 ml microcentrifuge tubes. Proteins (SMT3-PLIN2-GFP or PLIN3-APPLE) in protein dissolving buffer (50 mM Tris-HCl, 150 mM NaCl, pH 7.4) were added in varying concentrations to a final volume of 60 µl. The mixture was vortexed gently three times and centrifuged at 1,000 × *g* for 10 seconds. Then the samples were incubated in the at 4ºC in the dark for 12 h. After incubation, the adiposomes were centrifuged at 21,130 × *g* for 5 min to allow removal of the underlying solution. The adiposomes were then resuspended in 30 µl of Buffer B and centrifuged again. The wash procedure was repeated three times to remove the nonspecifically bound proteins. Finally, the adiposomes were resuspended in 800 µl of protein dissolving buffer and vortexed for at least 15 seconds. Then the samples were distributed equally into wells of a 96-microwell plate (200 µl for each well) in three technical replicates. At the same time, proteins were diluted into different concentrations with dissolving buffer containing Triton X-100 as standards. The final volume of each tube was also 800 µl and the final concentration of Triton X-100 was 1%. The standards were also distributed into the same 96-microwell plate using the same procedure. The plate was centrifuged at 3,220 × *g* for 1 min to remove bubbles and analyzed using an EnSpire Multimode Plate Reader. After the temperature of the machine stabilized at 25ºC, the absorption at 600 nm was measured. Then, fluorescence intensity (FI) was measured with excitation and emission at 488 nm and 530 nm, respectively, for GPF or 550 nm and 580 nm, respectively, for APPLE. One standard curve and one saturation curve were recorded in each experiment. At least two independent experiments were conducted in each case.

**Saturation analysis**

Protein purity was estimated using SDS-PAGE gels stained overnight using a Colloidal Blue Staining Kit (Invitrogen) and densitometry conducted with ImageJ. The percent purity was used in protein concentration normalization. A standard curve of fluorescence versus protein concentration was constructed and later used to determine the amount of protein bound to adiposomes. Background fluorescence, as measured in buffer, was subtracted from standards. The experiments were conducted in adiposome preparations with a starting OD600 of 20. Upon completion of the experiment, prior to the fluorescence measurement, adiposome preparations were diluted 13.3-fold to get into the linear range of the assay. The OD600 values of the diluted samples were measured and the ratio of $OD600_{Measured}$ to $OD600_{Theoretical}$ was used to correct for losses during washing when calculating the concentration of bound protein. Finally, the data was analyzed by nonlinear regression of $Conc_{Bound}$ versus $Conc_{Total}$.

**Scatchard analysis**

Scatchard analysis used to calculate the binding affinity and the maximum saturation concentration of binding sites on adiposomes. A linear regression of the plot of bound/free (ordinate) against bound (abscissa) yielded the slope = $-K_d^{-1}$, where $K_d$ is the equilibrium dissociation constant and the abscissa intercept = $B_{max}$, the maximum saturation concentration of ligand binding sites. The regressions were plotted using Graphpad 7.0.

**Structural analysis of PLIN2 and PLIN3**

The structures of PLIN2 (Homo sapiens) and PLIN3 (Homo sapiens) were modeled by I-TASSER (Yang and Zhang, 2015). PLIN3 (Mus musculus) was used as the template for homology modeling since the C-terminal (191-437) of PLIN3 (Mus musculus) has been structurally characterized (PDB ID: 1SZI, http://www.rcsb.org/) (Hickenbottom et al., 2004). The sequences of PLIN2 and PLIN3 which exhibited the highest confidence values were predicted using Heliquest and the structural comparison was performed using PyMOL (Chong et al., 2011).

**Single site-directed mutation of PLIN2 and ATGL**

The cloning and mutagenesis primers designed by VectorNTI were listed in Table S1 and the constructed plasmids were listed in Table S2. C2C12/Huh7 cells were used as the templates to perform the reverse transcription of RNA. The reaction system and procedure are shown in supplementary data. 1 μl of enzyme Dpn1 was added to the vector mixture and incubated at 37ºC water bath for 1 h. The TOP10 competent cells were mixed with the vector after enzymatic reaction in the ice bath for 30 min. They underwent the heat stimulus (42ºC) for 90 seconds and then were placed on ice for 2 min. 500 µl of LB medium was mixed with them and the cells were revived at 37ºC for 45 min with 200 rpm shaking. 50-100 µl of bacterial cells were coated on the resistant plates and cultured. pET-28a-SMT3-N plasmid was obtained from Dr. Sarah Perret's Lab. All plasmids were sequenced to confirm successful mutagenesis.

**Gene overexpression and knockout**

Huh7 cells were cultured in confocal dishes. The vector was dispersed in 500 µl of Opti-MEM medium and 5 µl of Lipofectamine 2000 was also dispersed in 500 µl of Opti-MEM medium. The vector suspension and Lipofectamine 2000 suspension were then mixed gently and incubated for 15-20 min. Cells were detected using fluorescence microscopy after well cultured. C2C12 cells were digested by trypsin. The digested cells were collected and centrifuged at 500 × *g* for 5 min. The supernatant was removed and the cells were resuspended using NT buffer. Cells were mixed with vector and electroporated. Those cells were resuspended and cultured for fluorescence microscope observation.

CRISPR/Cas9 technique was used to knockout ATGL. The mRNA sequence and genomic sequence were introduced into VectorNTI database to design the target. The target genome was identified using Crispr tools and the primers with the highest score were selected. The test primer was designed by introducing genome sequence into VectorNTI and screened the sequences (500-550 bp) that covered target sequence. The pX260a vector (a gift from Prof. Feng Zhang), T4 ligase, T4 ligase buffer and annealing product were used to construct the vector. The product was introduced into TOP10 competent cells and the sequence was verified after culturing. Afterwards, the correct vectors were collected and transfected into C2C12 cells. The gene knockout cells were cultured and screened using 1 µg $ml^{-1}$ Puromycin for 2 weeks. The cells were further diluted to gradients and cultured for 2-3 weeks. Next monoclone was selected and cultured, to screen the positive cells by immunoblotting and PCR. The positive cells were cultured and fed with oleic acid (50-100 μM) for 12-24 h, and then they were observed using fluorescence microscope.

**Triacylglycerol hydrolase assay**

The reagents were prepared as previously published (Schweiger et al., 2014) with the following modifications. The protease inhibitor in Solution A was replaced with PMSF and the n-heptane in extraction solution I (methanol/chloroform/n-heptane = 10:9:7, v:v:v) was substituted with n-hexane. Brown adipose tissue (BAT) cytosol from a C57BL/6 mouse (Vital River Laboratories) was suspended in Solution A (0.25 M sucrose, 1 mM EDTA, 1 mM DTT and 0.5 mM PMSF), flash frozen in liquid nitrogen, and stored at -80ºC. The ATGL mutants were expressed in Transetta (DE3) following the method described above. The bacterial supernatant (lysate) was obtained by sonication on ice in 1.6 ml of Solution A per 800 ml bacteria. Sonication was conducted for 15 min (6 seconds on, 6 seconds off) with an output power of 35% and was followed by centrifugation at 21,130 × *g* for 10 min. 25 µl of Solution A was used as a control. The radioactive labeled adiposomes were prepared using 1,100 µg DOPC, 340 µg liver PI, 520 µg DOPE (DOPC:liver PI:DOPE = 11:3:5, molar ratio) and 5 µl of TAG, mixed with 5 µl Triolein [9, 10-3H(N)] (0.5 µCi $µl^{-1}$, PerkinElmer). The solvents were dried using a stream of nitrogen. The adiposomes were produced using the method cited previously but were resuspended in Solution A in the last step. 25 µl of the bacterial supernatant was mixed with 25 µl of adiposome and the mixture was incubated for 1 h in at 37°C. The reaction was terminated by the addition of 650 µl extraction solution I followed by 200 µl of extraction solution II (0.1 M $K_2CO_3$, adjusting pH to 10.5 using boric acid) and vigorous votexing. Samples were centrifuged at 1,000 × *g* for 10 min to drive the proteins into the aqueous-organic boundary. 200 µl of the upper aqueous phase was transferred to a scintillation vial containing 1 ml of Optifluor liquid and its radioactivity was analyzed by gamma meter (PerkinElmer).

25 μl of the substrate was measured to determine specific substrate radioactivity. The lipase activity was calculated using the equation (Zimmermann et al., 2004):

$$nmol\ FA/_{mg\ protein} = \frac{(cpm_{Sample} - cpm_{Blank}) \times (V_1/V_2)}{\left(\frac{cpm_{Substrate}}{nmol\ FA}\right) \times mg\ protein \times 0.715t}$$

where $V_1$ is the total volume of upper water phase (2.45 ml); $V_2$ is the volume measured in the scintillation counter (0.2 ml); and t is the incubation time (h). The mass of protein is estimated by analyzing stained gels using ImageJ.

**Figures and Legends**

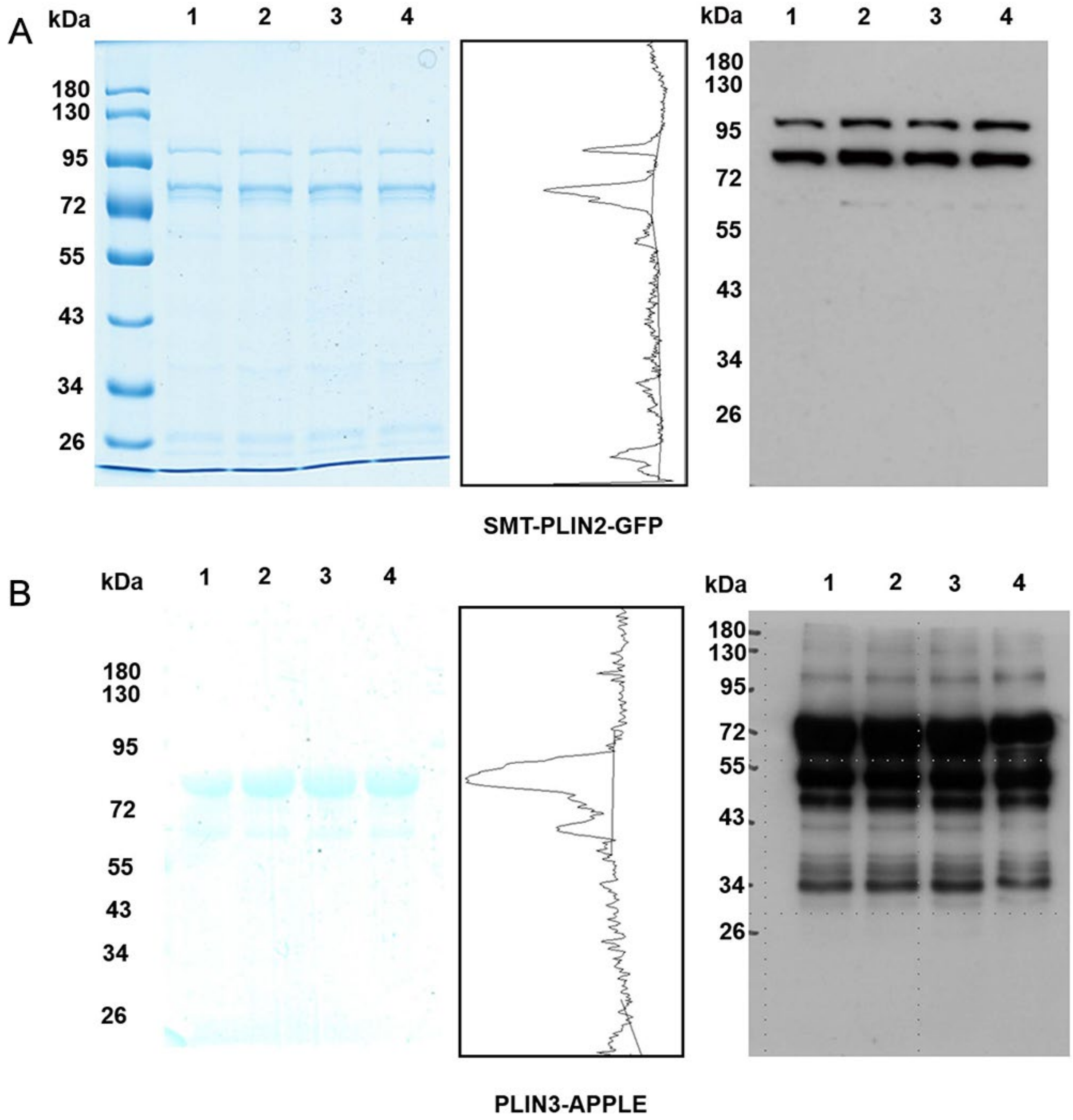


**Figure S1. The purification of recombinant PLIN2 and PLIN3.**

(A) SMT3-PLIN2-GFP and (B) PLIN3-APPLE were purified using our published method. Both proteins were identified using Colloidal Blue Staining and Western blot. The numbers on the top of figures represent four repeats. The Western blot was performed using anti-PLIN2 antibody for SMT3-PLIN2-GFP, and anti-PLIN3 antibody for

PLIN3-APPLE. The Colloidal Blue Staining figures were scanned using ImageJ to calculate the purity of proteins: the purity of SMT3-PLIN2-GFP and PLIN3-APPLE was 59% and 86%.

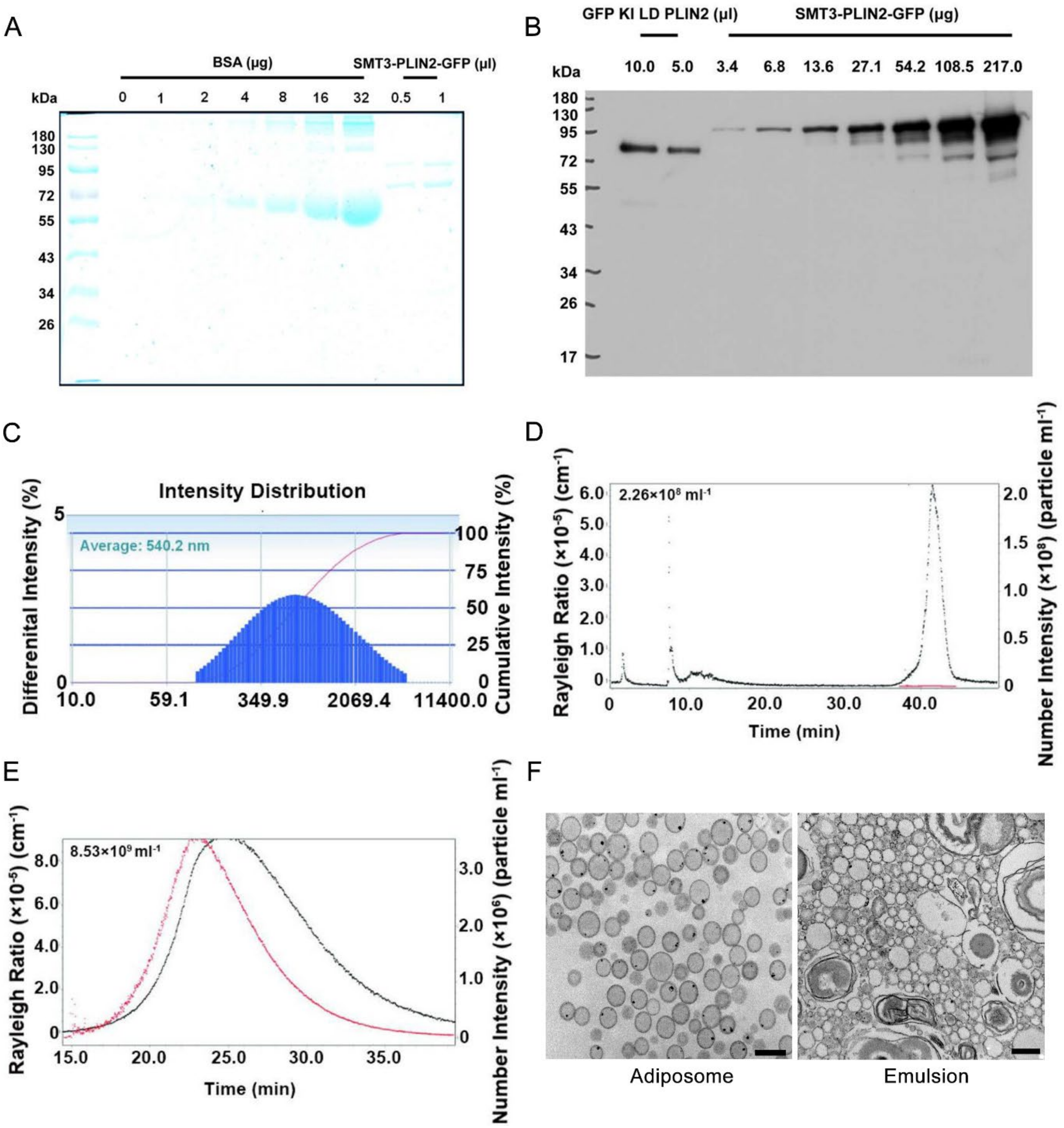


**Figure S2. The density of PLIN2 on lipid droplets and adiposomes.**

SMT3-PLIN2-GFP was expressed in bacteria and purified by affinity chromatography. (A) An aliquot of the SMT3-

PLIN2-GFP preparation was electrophoresed together with different amounts of BSA, and the gel was stained with Colloidal Blue Staining Kit overnight. ImageJ was used for quantification by densitometry to determine the original concentration of the SMT3-PLIN2-GFP preparation. (B) A dilution series of recombinant PLIN2 was compared to the PLIN2 content of lipid droplets purified from PLIN2–GFP knock-in C2C12 cells by Western blot with anti-PLIN2 antibody. ImageJ was used for quantification. The amount of endogenous PLIN2 was corrected for the molecular weight difference to SMT3-PLIN2-GFP. (C) Diameter of lipid droplets was 540.2 nm determined by DLS. Polydispersity Index = 0.341. (D) The concentration of purified lipid droplets was approximately $2.26 \times 10^8$ per ml measured by FFF-MALS system. (E) OD600 of the adiposome preparation was adjusted to 20 which had a concentration of approximately $8.53 \times 10^9$ per ml, measured with the FFF-MALS system. The red scatters and black scatters in (D) and (E) represented the number density of peak and LS 90.00º , respectively. (F) Morphology of adiposomes and lipid emulsions, which were observed using TEM by ultrathin section. Scale bar, 500 nm.

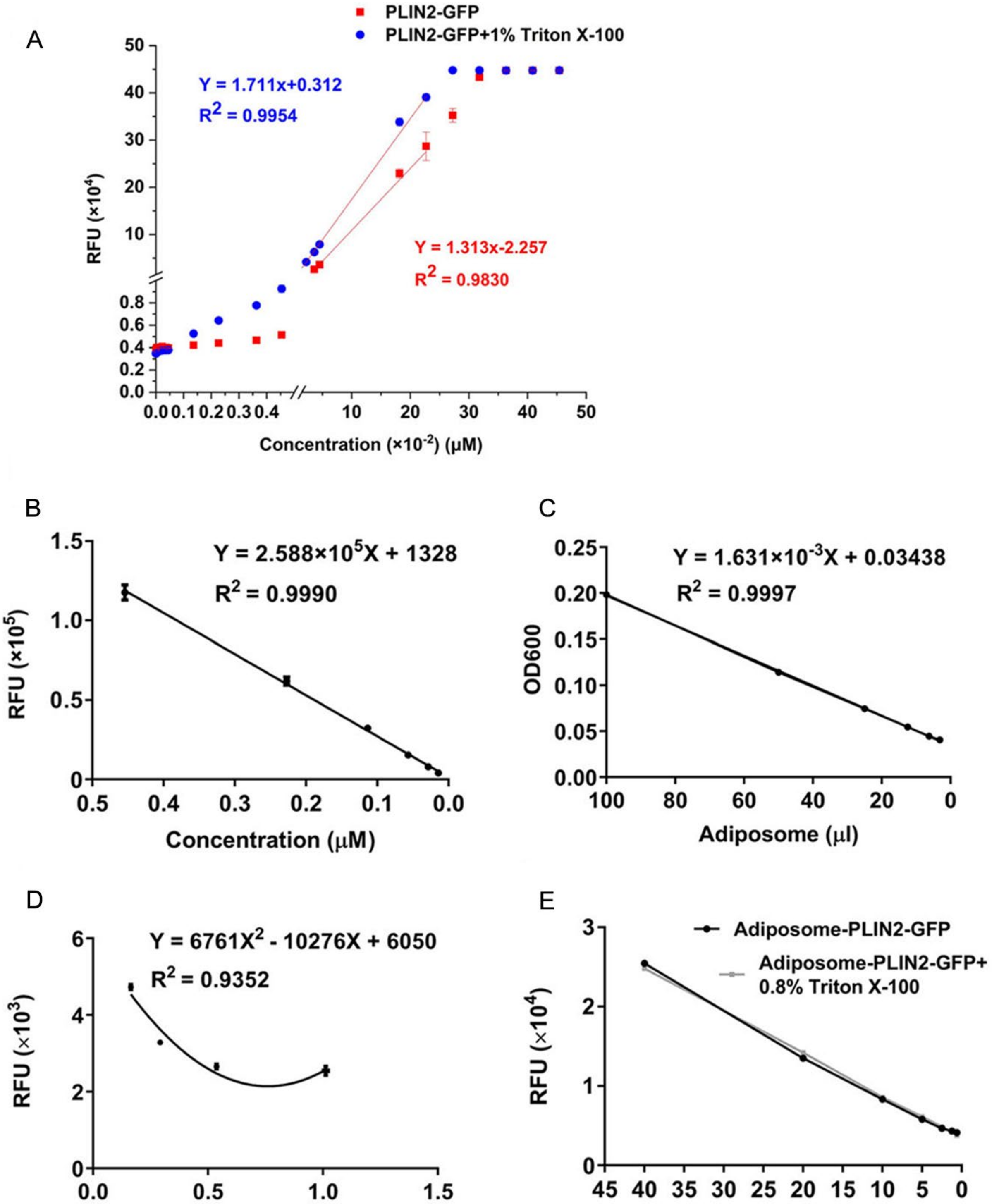


**Figure S3. The detection range of multi-scan spectrum with protein samples in a series of concentrations.**

(A) A series of protein samples of SMT3-PLIN2-GMP with concentrations range from 0 μM to 0.45 μM, from 0 μM to 0.045 μM, from 0 μM to 0.0045 μM, and from 0 μM to 0.0008 μM were scanned to obtain their fluorescence intensity, measured in Relative Fluorescence Units (RFU). A linear correlation between protein concentration and RFU was found from 0.005 μM to 0.25 μM. Protein in buffer containing 1% Triton X-100, to disperse potential protein aggregates, had slightly stronger fluorescence intensity. (B) There was a linear correlation between the concentration of protein and fluorescence intensity when the protein was diluted. (C) OD600 decreased linearly with decreasing adiposome concentration. Therefore, the OD600 value could be used to unify the amount of adiposomes for each assay for determining the protein-adiposome binding saturation curves. (D) By diluting adiposomes 2-fold, 4-fold, 8-fold, and 16-fold, a non-linear correlation was found between RFU and OD600. This is used to unify the adiposome loading. By creating the relation between RFU and OD600, the RFU of adiposome could be correctly identified after diluting for fluorescence scan. This indicated that OD600 could only be used for unifying the initial adiposome loading, rather than be used as the concentration value of adiposomes for the fluorescence scan. (E) After incubation with SMT3-PLIN2-GFP (at a final concentration of 0.3 μM), adiposomes were diluted by 2-fold, 4-fold, 8-fold, 16-fold, 32-fold, and 64-fold. Fluorescence intensity and adiposome quantity showed a linear relationship in the presence or absence of Triton X-100. This indicated that the proteins did not aggregate when incubated with adiposomes. These experiments were conducted to screen the linear range of fluorescence intensity of proteins when they were incubated with adiposomes, and therefore underlay the saturation curve depicting of protein binding.

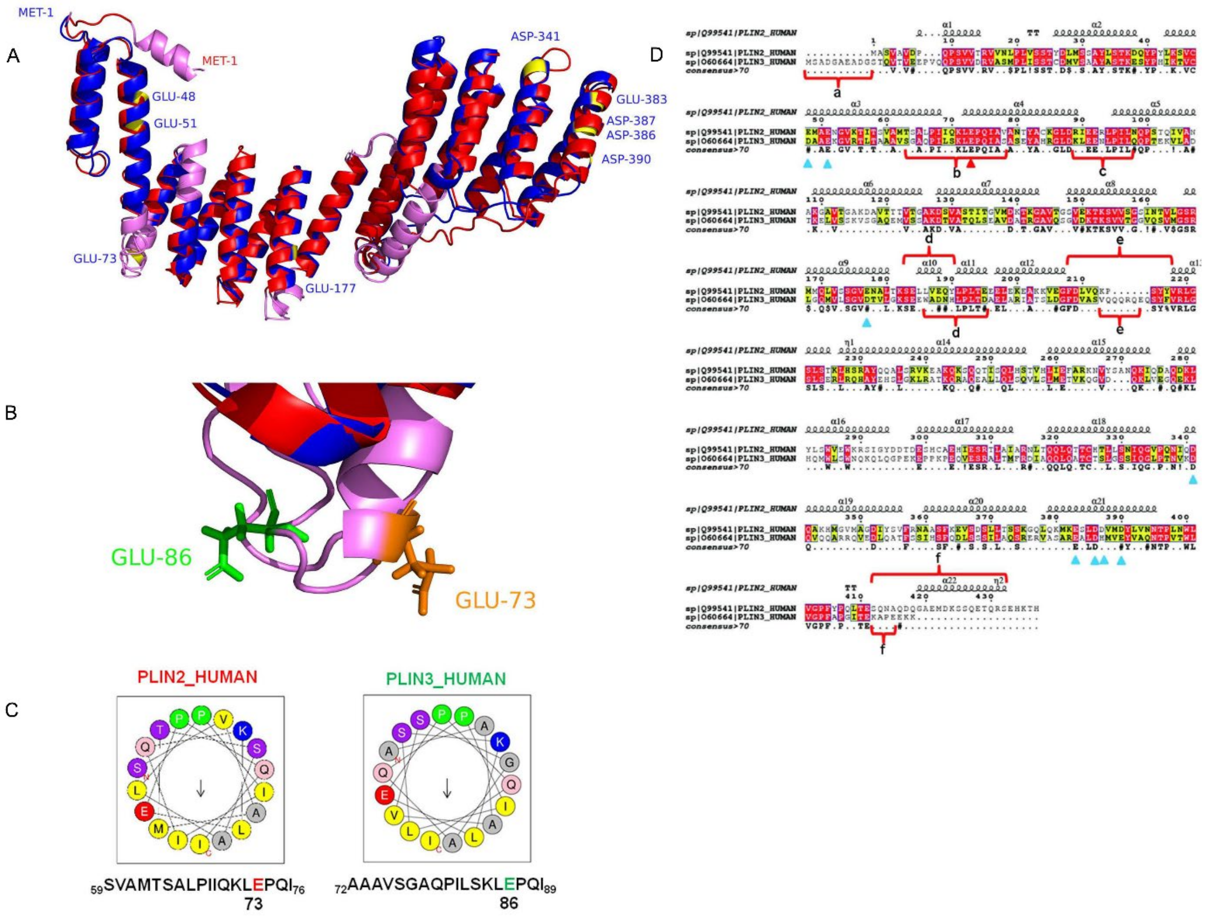


**Figure S4. The location of negatively charged amino acids in amphipathic α-helixes of PLIN2.**

The secondary structures of PLIN2 and PLIN3 were predicted by Phyre2.0 and the sequence of PLIN2 was visualized by IBS. The α-helixes of PLIN2 were predicted by HeliQuest and compared with the structure of corresponding sequences in PLIN3. The secondary structure of the sequence in the red box was different between two proteins and the location of glutamic acid was also different.

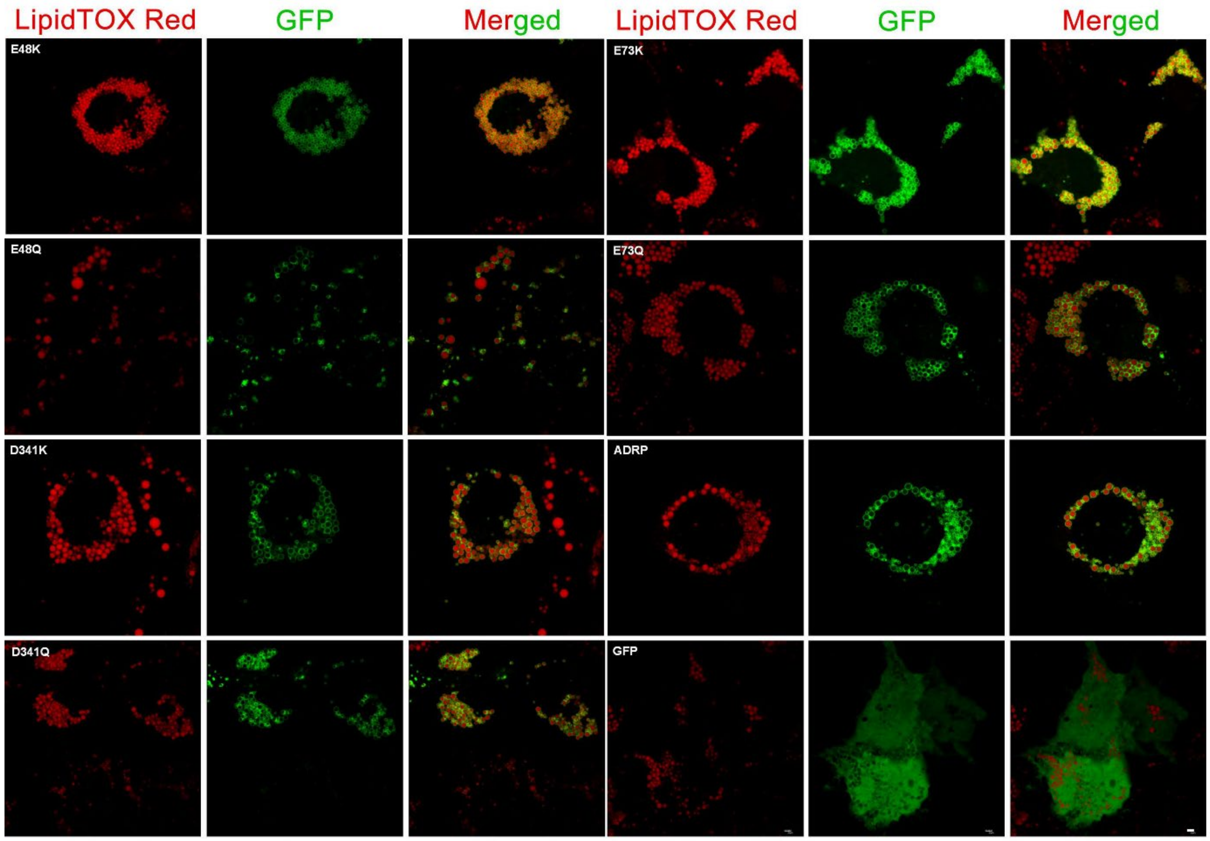


**Figure S5. The localization of GFP-tagged PLIN2 mutants on lipid droplet surface.**

PLIN2 mutants were overexpressed in Huh7 cells. Lipid droplets were stained with LipidTOX Red. Fluorescence images were obtained using a confocal microscope. Scale bar, 5 μm.

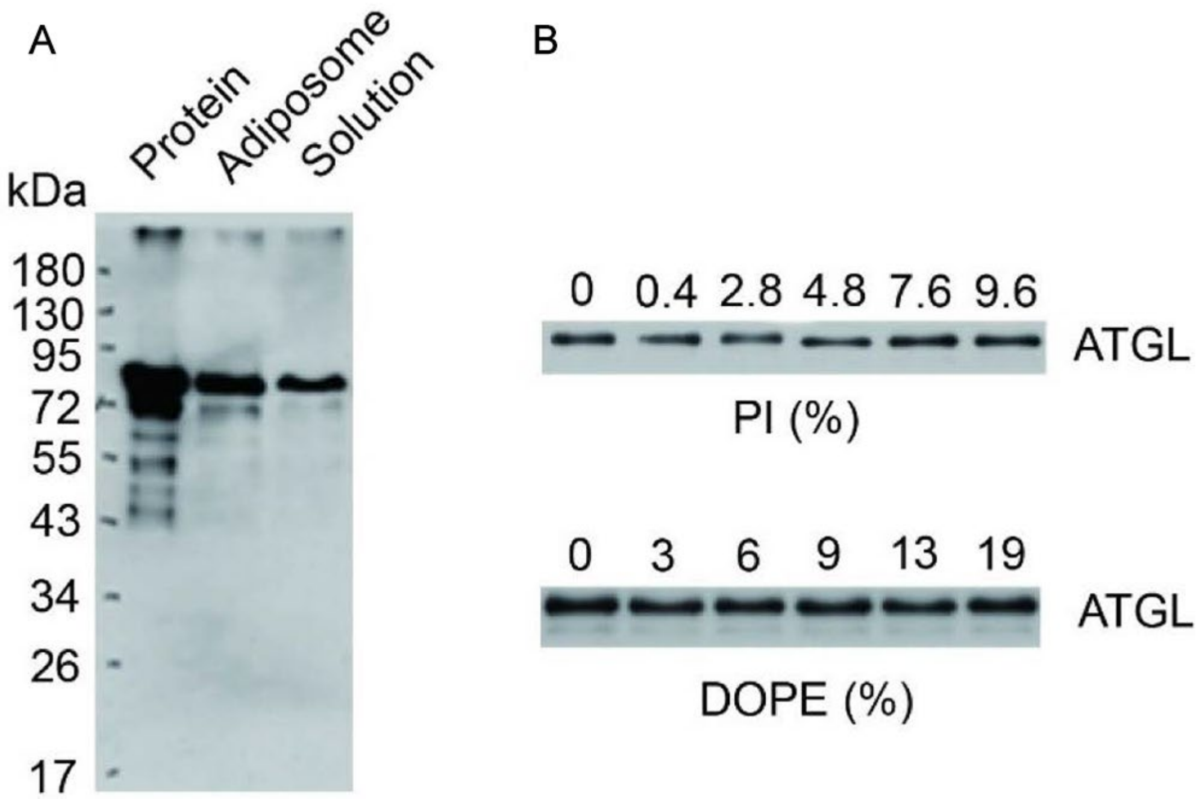


**Figure S6. The targeting of ATGL to adiposomes.**

12 μg of ATGL was incubated with 100 μl adiposomes for 1 h at room temperature. The adiposomes were centrifuged and washed three times. (A) The samples were analyzed by Western blot with anti-ATGL antibody, which showed that ATGL was able to target to adiposomes. (B) A series of adiposomes prepared using different concentrations of PI or DOPE were incubated with ATGL. There were no significant changes in ATGL targeting with the increasing ratio of PI or DOPE in adiposomes.

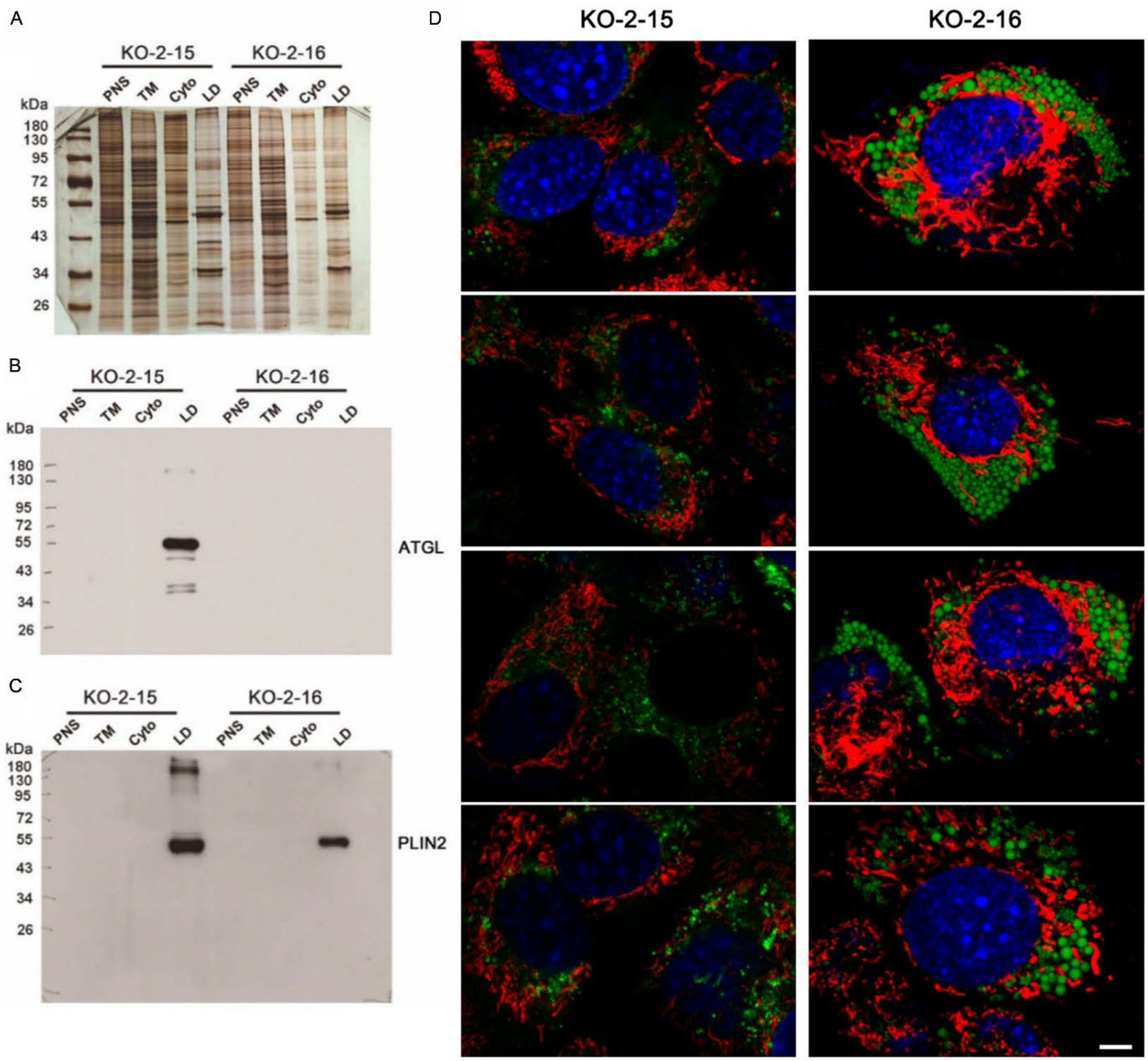


**Figure S7. The knockout of ATGL in C2C12 cells.**

The sequencing results of the monoclonal KO-2-15 and KO-2-16 cell lines showed that there was no deletion or insertion in the ATGL locus of KO-2-15 while there was a single base deletion in KO-2-16 causing frameshift in the ATGL gene. (A) Lipid droplets were purified from these two clones and the protein profiles were analyzed by silver staining. (B) The cell fractions were further analyzed by Western blot with anti-ATGL antibody. There was no ATGL expression in the KO-2-16 cell line. (C) The expression of PLIN2 was decreased in the ATGL knock-out cells as detected by anti-PLIN2 antibody. (D) Cells grown overnight in confocal dishes were stained using LipidTOX Green

(LDs, green) and MitoTracker Red (mitochondria, red) and Hoechst (nuclei, blue). The cells were examined using an Olympus FV1000 confocal microscope. Both images were obtained using the same exposure time and microscopy settings. The images showed that lipid droplets and mitochondria were larger in ATGL knock-out cells. Scale bar, 5 μm.

**Table S1. The primers used in this study.**

| Primer | Sequence (5'-3') |
|---|---|
| | hADRP amphiphilic α-helix acidic amino acid mutants |
| E48K F | TACCTGAA GTCTGTGTGT AAGATGGCAG AGAAC |
| E48K R | CTTACACA CAGACTTCAG GTAGGGATAC TGGTC |
| E48Q F | TACCTGAA GTCTGTGTGT CAGATGGCAG AGAAC |
| E48Q R | CTGACACA CAGACTTCAG GTAGGGATAC TGGTC |
| E51K F | TCTGTGTG TGAGATGGCA AAGAACGGTG TGAAG |
| E51K R | CTTTGCCA TCTCACACAC AGACTTCAGG TAGGG |
| E51Q F | TCTGTGTG TGAGATGGCA CAGAACGGTG TGAAG |
| E51Q R | CTGTGCCA TCTCACACAC AGACTTCAGG TAGGG |
| E73K F | CCCATCAT CCAGAAGCTA AAGCCGCAAA TTGCA |
| E73K R | CTTTAGCT TCTGGATGAT GGGCAGAGCA CTGGT |
| E73Q F | CCCATCAT CCAGAAGCTA CAGCCGCAAA TTGCA |
| E73Q R | CTGTAGCT TCTGGATGAT GGGCAGAGCA CTGGT |
| E177K F | CTCGTGAG CAGTGGCGTA AAAAATGCAC TCACC |
| E177K R | TTTTACGC CACTGCTCAC GAGCTGCATC ATCCG |

| | |
|---|---|
| E177Q F | CTCGTGAG CAGTGGCGTA CAAAATGCAC TCACC |
| E177Q R | TTGTACGC CACTGCTCAC GAGCTGCATC ATCCG |
| D341K F | GTACCACA GAACATCCAA AAACAAGCCA AGCAC |
| D341K R | TTTTTGGA TGTTCTGTGG TACACCTTGG ATGTT |
| D341Q F | GTACCACA GAACATCCAA AACCAAGCCA AGCAC |
| D341Q R | GTTTTGGA TGTTCTGTGG TACACCTTGG ATGTT |
| E383K F | CAGCTGCA GAAAATGAAG AAATCTTTAG ATGAC |
| E383K R | TTTCTTCA TTTTCTGCAG CTGCCCCTTG CTAGA |
| E383Q F | CAGCTGCA GAAAATGAAG CAATCTTTAG ATGAC |
| E383Q R | TTGCTTCA TTTTCTGCAG CTGCCCCTTG CTAGA |
| E386K F | AAAATGAA GGAATCTTTA AAAGACGTGA TGGAT |
| E386K R | TTTTAAAG ATTCCTTCAT TTTCTGCAGC TGCCC |
| E386N F | AAAATGAA GGAATCTTTA AACGACGTGA TGGAT |
| E386N R | GTTTAAAG ATTCCTTCAT TTTCTGCAGC TGCCC |
| E387K F | ATGAAGGA ATCTTTAGAT AAAGTGATGG ATTAT |
| E387K R | TTTATCTA AAGATTCCTT CATTTTCTGC AGCTG |
| E387N F | ATGAAGGA ATCTTTAGAT AACGTGATGG ATTAT |
| E387N R | GTTATCTA AAGATTCCTT CATTTTCTGC AGCTG |
| E390K F | TCTTTAGA TGACGTGATG AAATATCTTG TTAAC |
| E390K R | TTTCATCA CGTCATCTAA AGATTCCTTC ATTTT |
| E390N F | TCTTTAGA TGACGTGATG AACTATCTTG TTAAC |
| E390N R | GTTCATCA CGTCATCTAA AGATTCCTTC ATTTT |
| | mATGL phosphorylation site mutants |
| S47A F | CACATCTACG GAGCCGCGGC AGGGGCG |
| S47A R | CGCGGCTCCGT AGATGTGAGT GGCGTTG |
| S47D F | CACATCTACG GAGCCGACGC AGGGGCGC |
| S47D R | GTCGGCTCCGT AGATGTGAGT GGCGTTG |

| | |
|---|---|
| S87A F | CCTCTGCATC CCGCGTTCAA CCTGGT |
| S87A R | CGCGGGATGC AGAGGACCCA GGAACC |
| S87D F | CCTCTGCATC CCGACTTCAA CCTGGT |
| S87D R | GTCGGGATGC AGAGGACCCA GGAACC |
| T101A F | TGTCTACTAA AGGCGCTGCC TGCTGA |
| T101A R | CGCCTTTAGT AGACAGCCAC GGATG |
| T101D F | TGTCTACTAA AGGACCTGCC TGCTGA |
| T101D F | GTCCTTTAGT AGACAGCCAC GGATG |
| T210A F | CGCGTCACCA ACGCGAGCAT CCAGTT |
| T210A R | CGCGTTGGTGA CGCGAAGCT CGTGGA |
| T210D F | CGCGTCACCA ACGACAGCAT CCAGTT |
| T210D R | GTCGTTGGTGA CGCGAAGCT CGTGGA |
| T372A F | ATGAAAGAGC AGGCGGGTAG CATCT |
| T372A R | CGCCTGCTCT TTCATCCACC GGATA |
| T372D F | ATGAAAGAGC AGGACGGTAG CATCT |
| T372D R | GTCCTGCTCT TTCATCCACC GGATA |
| S393A F | GACCATCTGC CTGCGAGACT GTCTGA |
| S393A R | CGCAGGCAGA TGGTCACCCA ATTTC |
| S393D F | GACCATCTGC CTGACAGACT GTCTGA |
| S393D R | GTCAGGCAGA TGGTCACCCA ATTTC |
| Y378A F | AGCATCTGCC AGGCGCTGGT GATGA |
| Y378A R | CGCCTGGCAG ATGCTACCCG TCTGCT |
| Y378D F | AGCATCTGCC AGGACCTGGT GATGA |
| Y378D R | GTCCTGGCAG ATGCTACCCG TCTGCT |
| S396A F | CCTTCCAGAC TGGCGGAGCA GGTGGA |
| S396A R | CGCCAGTCTG GAAGGCAGAT GGTCA |
| S396D F | CCTTCCAGAC TGGACGAGCA GGTGGA |

| | |
|---|---|
| S396D R | GTCCAGTCTG GAAGGCAGAT GGTCA |
| S406A F | CTGCGACGTG CCCAGGCGCT GCCCTCTG |
| S406A R | CGCCTGGGCA CGTCGCAGTT CCACCTGC |
| S406D F | CTGCGACGTG CCCAGGACCT GCCCTCTG |
| S406D R | GTCCTGGGCA CGTCGCAGTT CCACCTGC |
| S430A F | GTACGAAACA ACCTCGCGCT GGGGGACG |
| S430A R | CGCGAGGTTG TTTCGTACCC AGTTGGGT |
| S430D F | GTACGAAACA ACCTCGACCT GGGGGACG |
| S430D R | GTCGAGGTTG TTTCGTACCC AGTTGGGT |
| | mATGL Knock-out |
| ATGL (mouse) KO Target-1 F | CACCGCGGGGTCTACCACATTGGCG |
| ATGL (mouse) KO Target-1 R | AAACCGCCAATGTGGTAGACCCCGC |
| ATGL (mouse) KO Target-2 F | CACCGATTGCCATGAGCGCGCCAA |
| ATGL (mouse) KO Target-2 R | AAACTTGGCGCGCTCATGGCAATC |
| ATGL KO Target-1 Test F | CCGGCGGAGACCCCAAGGTA |
| ATGL KO Target-1 Test R | ACAAACCAGAACCCGCCCCG |
| ATGL KO Target-2 Test F | GTCTTCACAGTCAATGGAAT |
| ATGL KO Target-2 Test R | AGGGTAGGAGGAATGAGGCC |
| | SMT3-hADRP-GFP fusion proteins |
| A1 GFP-ADRP-GFP-F | CCGGAATTCATGGCATCCGTTGCAGTTG |
| A2 | TCCTCGCCCTTGCTCACCATATGAGTTTTATGCTCA GATC |
| B1 | GATCTGAGCATAAAACTCATATGGTGAGCAAGGGC GAGGA |
| B2 pET-28a-EGFP-C-R | CCGCTCGAGTTACTTGTACAGCTCGTCCATGC |
| | hTIP47-APPLE fusion proteins |
| A1 APPLE-TIP47 F | CCGGAATTCATGTCTGCCGACGGGGCAGA |

| | |
|---|---|
| A2 | TCCTCGCCCTTGCTCACCATCTTCTTCTCCTCCGGGGCTT |
| B1 | AAGCCCCGGAGGAGAAGAAGATGGTGAGCAAGGGCGAGGA |
| APPLE R | CCGCTCGAGTTACTTGTACAGCTCGTCCA |

**Table S2. The plasmids used in this study.**

| Plasmid | |
|---|---|
| hPLIN2 mutant using pET-28a-SMT3-N, expressed in Transetta(DE3) | SMT3-hADRP-GFP E48K |
| | SMT3-hADRP-GFP E48Q |
| | SMT3-hADRP-GFP E51K |
| | SMT3-hADRP-GFP E51Q |
| | SMT3-hADRP-GFP E73K |
| | SMT3-hADRP-GFP E73Q |
| | SMT3-hADRP-GFP E177K |
| | SMT3-hADRP-GFP E177Q |
| | SMT3-hADRP-GFP D341K |
| | SMT3-hADRP-GFP D341Q |
| | SMT3-hADRP-GFP E383K |
| | SMT3-hADRP-GFP E383Q |
| | SMT3-hADRP-GFP E386K |

| | |
|---|---|
| | SMT3-hADRP-GFP E386N |
| | SMT3-hADRP-GFP E387K |
| | SMT3-hADRP-GFP E387N |
| | SMT3-hADRP-GFP E390K |
| | SMT3-hADRP-GFP E390N |
| hPLIN2 mutant using pEGFP-N1, expressed in Huh7 cells | hADRP-GFP E48K |
| | hADRP-GFP E48Q |
| | hADRP-GFP E51K |
| | hADRP-GFP E51Q |
| | hADRP-GFP E73K |
| | hADRP-GFP E73Q |
| | hADRP-GFP E177K |
| | hADRP-GFP E177Q |
| | hADRP-GFP D341K |
| | hADRP-GFP D341Q |
| | hADRP-GFP E383K |
| | hADRP-GFP E383Q |
| | hADRP-GFP E386K |
| | hADRP-GFP E386N |
| | hADRP-GFP E387K |

| | |
|---|---|
| | hADRP-GFP E387N |
| | hADRP-GFP E390K |
| | hADRP-GFP E390N |
| ATGL mutant using pFlag-CMV4, expressed in C2C12 cells | mATGL S47A |
| | mATGL S47D |
| | mATGL S87A |
| | mATGL S87D |
| | mATGL T101A |
| | mATGL T101D |
| | mATGL T210A |
| | mATGL T210D |
| | mATGL T372A |
| | mATGL T372D |
| | mATGL S393A |
| | mATGL S393D |
| | mATGL Y378A |
| | mATGL Y378D |
| | mATGL S396A |
| | mATGL S396D |
| | mATGL S406A |

| |
|---|
| mATGL S406D |
| mATGL S430A |
| mATGL S430D |

### References


Chong, B.M., Russell, T.D., Schaack, J., Orlicky, D.J., Reigan, P., Ladinsky, M., and McManaman, J.L. (2011). The adipophilin C terminus is a self-folding membrane-binding domain that is important for milk lipid secretion. J Biol Chem *286*, 23254-23265.

Hickenbottom, S.J., Kimmel, A.R., Londos, C., and Hurley, J.H. (2004). Structure of a lipid droplet protein: the PAT family member TIP47. Structure *12*, 1199-1207.

Schweiger, M., Eichmann, T.O., Taschler, U., Zimmermann, R., Zechner, R., and Lass, A. (2014). Chapter Ten - Measurement of lipolysis. In Methods Enzymol, O.A. MacDougald, ed. (Academic Press), pp. 171-193.

Wang, Y., Zhou, X.-M., Ma, X., Du, Y., Zheng, L., and Liu, P. (2016). Construction of nanodroplet/adiposome and artificial lipid droplets. ACS Nano *10*, 3312-3322.

Yang, J., and Zhang, Y. (2015). I-TASSER server: new development for protein structure and function predictions. Nucleic Acids Res *43*, W174-W181.

Zimmermann, R., Strauss, J.G., Haemmerle, G., Schoiswohl, G., Birner-Gruenberger, R., Riederer, M., Lass, A., Neuberger, G., Eisenhaber, F., Hermetter, A., *et al.* (2004). Fat mobilization in adipose tissue is promoted by adipose triglyceride lipase. Science *306*, 1383-1386.
